# Divergent roles for caspase-8 and MLKL in high-fat diet induced obesity and NAFLD in mice

**DOI:** 10.1101/2023.03.14.532682

**Authors:** Hazel Tye, Stephanie A. Conos, Tirta M. Djajawi, Nasteho Abdoulkader, Isabella Y. Kong, Helene L. Kammoun, Vinod K. Narayana, Mary Speir, Timothy A. Gottschalk, Jack Emery, Daniel S. Simpson, Cathrine Hall, Angelina J. Vince, Sophia Russo, Rhiannan Crawley, Maryam Rashidi, Joanne M. Hildebrand, James M. Murphy, Lachlan Whitehead, David P. De Souza, Seth L. Masters, Edwin D. Hawkins, Andrew J. Murphy, James E. Vince, Kate E. Lawlor

## Abstract

Cell death frequently occurs in the pathogenesis of obesity and non-alcoholic fatty liver disease (NAFLD). However, the exact contribution of core cell death machinery to disease manifestations remains ill-defined. Here, we show *via* the direct comparison of mice genetically deficient in apoptotic caspase-8 in myeloid cells, or the essential necroptotic regulators, Receptor-interacting protein kinase-3 (RIPK3) and Mixed lineage kinase domain-like (MLKL), that RIPK3-caspase-8 signaling regulates macrophage inflammatory responses and drives adipose tissue inflammation and NAFLD upon high-fat diet feeding. In contrast, MLKL, divergent to RIPK3, contributes to both obesity and NAFLD in a manner largely independent of inflammation. We also uncover that MLKL regulates the expression of molecules involved in lipid uptake, transport and metabolism and, congruent with this, we discover a shift in the hepatic lipidome upon MLKL deletion. Collectively, these findings highlight MLKL as an attractive therapeutic target to combat the growing obesity pandemic and metabolic disease.

## Introduction

Obesity is a global pandemic associated with the consumption of a Western diet rich in saturated fats and refined carbohydrate, and a sedentary lifestyle. Diet-induced obesity leads to increased circulating free fatty acids, insulin and glucose that trigger “low grade” inflammation that is causally-related to inflammatory macrophage expansion within hypertrophic adipose tissue^1^. This state of chronic metabolic inflammation predisposes obese individuals to multi-organ insulin resistance and co-morbidities, such as Non-Alcoholic Fatty Liver disease (NAFLD). Evidence implicates gut dysbiosis and a “leaky gut” in systemic inflammation *via* the release of damaging microbial products (*e.g*., lipopolysaccharide)^2^ that act synergistically with excess dietary metabolites, such as saturated fatty acids, to potentiate inflammatory signaling and cytokine/chemokine production^3^. Obesity-induced inflammation is also inextricably linked to the activation and assembly of the NOD-like receptor protein 3 (NLRP3) inflammasome through the sensing of various metabolic DAMPs (*e.g*., palmitic acid, cholesterol crystals)^4^. NLPR3 inflammasome-associated caspase-1 subsequently cleaves and activates IL-1β, as well as the pyroptotic effector gasdermin D (GSDMD)^5,6^. Intriguingly, despite clear evidence that NLRP3 inflammasome and IL-1β activity drive obesity and NAFLD^7–10^, whether pyroptosis, or other modes of programmed cell death, such as apoptosis and necroptosis, facilitate the demise of key cell types in tissues remains ambiguous.

Targeting the extensive hepatocyte death observed in NAFLD progression is thought to be attractive to prevent subsequent liver pathologies. Hepatocyte death has been widely attributed to extrinsic death receptor-mediated apoptotic caspase-8 activity. In this scenario, formation of the pro-apoptotic complex II, consisting of Receptor-interacting protein kinase-1 (RIPK1)-FADD-caspase-8 complex, or the related ripoptosome complex (RIPK1-RIPK3-FADD-caspase-8), may be triggered upon TNFR or TLR ligation, when pro-survival signaling is compromised (*e.g*., upon IAP loss)^11^. Fitting with a key role for apoptotic caspase-8 in NAFLD, deletion of caspase-8 in hepatocytes protected mice from Methionine-choline deficient (MCD)-diet induced liver injury and inflammation^12^, whilst pancaspase inhibitor emricasan (IDN-6556) attenuated liver injury, inflammation and fibrosis from high-fat diet (HFD) intake^13^. Conflictingly, caspase-8 deletion in mouse liver parenchymal cells exacerbated MCD-diet induced liver damage^14^, while emricasan underperformed in Phase II clinical trials and sometimes worsened progression of Non-alcoholic Steatohepatitis (NASH)^15^, suggesting that a form of caspase-independent cell death is triggered.

Necroptosis is a lytic form of cell death that is unleashed in the absence of caspase-8 activity and is critically dependent on RIPK1/3 kinase activity and the pseudokinase Mixed lineage kinase domain-like (MLKL). Formation of the necrosome triggers RIPK3-mediated phosphorylation of MLKL, prompting a conformational change that allows the N-terminal four-helical bundle domain to associate with membranes, causing cell lysis^16–19^. Whilst the role of upstream kinases RIPK1 and RIPK3 in obesity and NAFLD remain debatable^20–26^, there are numerous reports that MLKL deficiency protects mice from NAFLD^27–29^. As RIPK3 can regulate the activity of both caspase-8 and MLKL activity^30,31^, direct comparisons of mutant animals in the same obesity and NAFLD model is required to define their pathogenic roles and divergent activities.

Significant plasticity has been shown between apoptotic, pyroptotic and necroptotic cell death signaling pathways^32–36^. Moreover, crosstalk between intrinsic apoptosis, extrinsic apoptosis and necroptosis with NLRP3 inflammasome and IL-1 β activation has been documented in innate immune cells^31,37–43^ and *in vivo* in models of inflammatory disease and infection^31,44–49^. As both caspase-8 and MLKL signaling can be triggered by dietary metabolite excess, it remains to be seen whether either of these cell death modes act upstream of NLRP3, or act in parallel, to drive distinct aspects of pathology. These studies will be vital to the advancement of therapies in the area and for predicting disease outcomes. Here, we directly contrast the contribution of RIPK3, caspase-8 and MLKL signaling to inflammation and obesity and reveal that caspase-8 plays a role in limiting pathogenic inflammation to saturated fatty acids, whilst MLKL uniquely regulates obesity and NAFLD *via* noncanonical actions on lipid metabolism.

## Results

### Caspase-8 contributes to inflammasome priming, IL-1β activation and cell death upon LPS and palmitate exposure

To confirm the role of the NLRP3 inflammasome in IL-1β activation in macrophages in response to saturated fatty acid palmitate^50^, we examined inflammatory responses in LPS-primed wildtype (WT), NLRP3-deficient (*Nlrp3^-/-^*) and caspase-1-deficient (*Casp1^-/-^*) bone marrow-derived macrophages (BMDMs). As expected, NLRP3 or caspase-1 deficiency almost completely blocked caspase-dependent cleavage and activation of IL-1β in response to increasing concentrations of palmitate conjugated to bovine serum albumin (PA-BSA) (**Figures 1A, 1B and S1A**), with residual bioactive IL-1β likely due to caspase-8 activation (marked by the presence of p43 and p18 cleavage products) (**Figures 1F and S1J**). In line with previous reports^3^, we also observed that LPS and palmitate amplify inflammatory responses, shown by heightened TNF secretion (a marker of inflammasome priming) in WT and inflammasome-deficient BMDMs (**Figures 1C and S1B**). Intriguingly, cell death at 18-20 hours post LPS and palmitate exposure was not perturbed in *Nlrp3^-/-^* or *Casp1^-/-^* BMDMs (**Figures 1D and S1C**), whereas the canonical NLRP3 inflammasome stimulus nigericin failed to induce both IL-1β and pyroptotic cell death in the absence of NLRP3 or caspase-1 (**Figures S1D-S1F**). These results suggest that while palmitate induces NLRP3 inflammasome activation in LPS-primed macrophages, ultimately, cellular demise occurs independent of pyroptosis.

**Figure 1.**
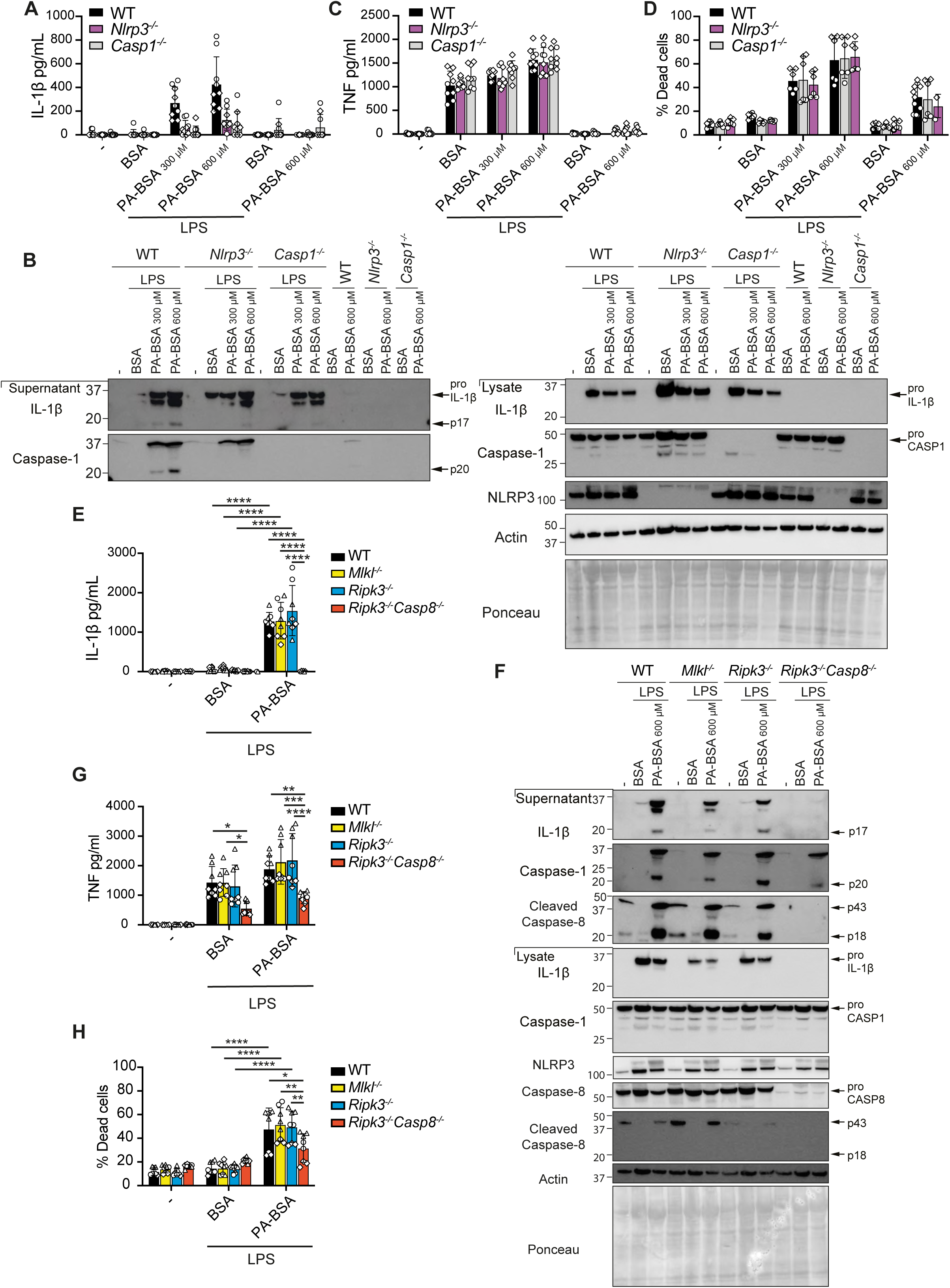
Caspase-8 activity contributes to LPS and palmitate-induced NLRP3 inflammasome priming, IL-1β activation and macrophage cell death. **(A-D)** WT, *Nlrp3^-/-^* and *Casp1^-/-^* BMDMs were pre-treated with or without LPS (50 ng/ml) for 3 h and treated with 300-600 μM palmitate conjugated to BSA (PA-BSA) or BSA alone (equivalent to 600 μM BSA amount) for a further 18 h. **(A)** IL-1β and **(C)** TNF levels were measured in cell supernatants by ELISA. (n = 4-5 replicates, 2 experiments). **(B)** Cell lysates and supernatants were analyzed by immunoblot for specified proteins. One of 2 experiments. **(D)** Cell viability was assessed by propidium iodide (PI) incorporation and flow cytometric analysis. n = 4-5 replicates from 2 independent experiments. **(E-H)** WT, *Mlkl^-/-^*, *Ripk3^-/-^* and *Ripk3^-/-^Casp8^-/-^* BMDMs were primed with or without LPS (50 ng/ml) for 3 h, as indicated, and treated with 600 μM PA-BSA or BSA alone, as indicated, for 18 h. **(E)** IL-1β and **(G)** TNF levels were measured in cell supernatants by ELISA. (n = 2-3 replicates from 3 experiments). **(F)** Cell lysates and supernatants were analyzed by immunoblot for specified proteins. One of 2 blots. **(H)** Cell viability was assessed by PI uptake and flow cytometric analysis. (n = 2-3 replicates, 3 experiments). For individual experiments, replicates are shown as different shapes. Data are the mean ± SD. *p < 0.05; **p < 0.01; ***p < 0.001; ****p < 0.0001. Two-way ANOVA followed by Tukey’s multiple comparisons test was applied. See related **Figure S1.**

Extrinsic apoptotic and necroptotic signaling have been shown to culminate in potassium ion efflux-dependent NLRP3 inflammasome activation^37^. Active caspase-8 can also directly cleave IL-1 β to its bioactive p17 form^30,51^. We therefore queried whether caspase-8 or MLKL signaling could trigger NLRP3 upon dietary stress. As caspase-8 deficiency induces lethal necroptotic signaling during embryogenesis^52^, we compared responses in mice lacking both necroptotic RIPK3 and caspase-8 (*Ripk3^-/-^Casp8^-/-^*) with RIPK3-deficient (*Ripk3^-/-^*) and MLKL-deficient (*Mlkl^-/-^*) mice. Interestingly, IL-1β activation was completely blocked in *Ripk3^-/-^ Casp8^-/-^* LPS-primed BMDMs after 18-20 hours of palmitate stimulation, while loss of MLKL or RIPK3 had no impact (**Figures 1E and 1F)**, suggesting that caspase-8 activity regulates NLRP3 inflammasome and IL-1 β activity. However, in line with previous reports of a transcriptional role for caspase-8 in inflammasome priming^53,54^, LPS-primed *Ripk3 Casp8* BMDMs exhibited reduced pro-IL-1β, NLRP3 and TNF levels compared with WT cells (**Figures 1F and 1G**) and, accordingly, blunted IL-1β secretion to nigericin (**Figures S1G and S1H**). Importantly, the presence of caspase-1 (p20) activity in LPS and PA-BSA treated *Ripk3^-/-^ Casp8^-/-^* BMDMs (**Figure 1F**), albeit reduced compared to WT, suggested that caspase-8 is not essential for palmitate-induced NLRP3 activation. Consistent with this, following Pam3Cys (TLR2) priming, which is less dependent on caspase-8-induced transcriptional^53^ and post-translational effects^55^, we observed that palmitate triggered relatively normal caspase-1 activity and appreciable IL-1β (p17) secretion in *Ripk3^-/-^ Casp8^-/-^* BMDMs (**Figure S1I-S1K**). Parallel analysis of cell death revealed that RIPK3 and MLKL were also not required for TLR and PA-BSA-induced cell death at 20 hours, whilst caspase-8 deficiency modestly reduced cell death in LPS-primed cells (**Figures 1F, 1H and S1J, S1L**). Collectively, this suggests that caspase-8 not only regulates inflammasome priming in macrophages but partly contributes to IL-1 β proteolysis and cell death.

### Caspase-8 activity in myeloid cells contributes to obesity-induced metabolic dysfunction

Based on the inflammatory effects of caspase-8 in macrophages *in vitro*, we next investigated how loss of caspase-8 in myeloid cells would impact obesity. As *Ripk3^-/-^ Casp8^-/-^* mice develop an autoimmune lymphoproliferative syndrome that precludes their long-term analysis in a high-fat diet (HFD) obesity model^52^, we examined mice lacking caspase-8 conditionally in myeloid cells (LysMcre transgene) on a RIPK3-deficient background (*Casp8^LysMcre^Ripk3^-/-^*)^41^. On a normal-chow diet (ND), *Casp8^LysMcre^Ripk3^-/-^* mice displayed no major difference in weights with ageing, apart from a minor yet significant reduction in % weight gain compared to *Casp8^lox/lox^Ripk3^-/-^* (RIPK3-deficient) and WT *Casp8^lox/lox^* control mice (**Figures 2A and S2A**). In contrast, over 25 weeks of HFD feeding, *Casp8^yMy^Ripk3^-/-^* mice gained significantly less weight than control mice, while RIPK3-deficient mice exhibited delayed weight gain (**Figures 2B and S2B**). This discrepancy in weight was not grossly attributable to altered food intake or fecal output (**Figure S2C**). After 25 weeks on ND, there were no differences in subcutaneous adipose tissue (SAT) and visceral adipose tissue (VAT) expansion, with a modest reduction in liver weights in both *Casp8^lox/lox^Ripk3^-/-^* and *Casp8^LysMcre^Ripk3*^-/-^ animals, compared to control mice (**Figures 2C and S2D, S2F**). Likewise, all HFD-fed mice displayed comparable enlargement of SAT and VAT (**Figures 2D and S2E, S2F**), and RIPK3-deficient mice, in fact, tended to be more obese. Impaired weight gain in HFD-fed *Casp8^LysMcre^Ripk3^-/-^* mice instead appeared attributable to reduced liver enlargement, suggesting that myeloid caspase-8 activity contributes to disease progression (**Figures 2D and S2F**). RIPK3 deficiency alone also appeared to confer some protection from liver enlargement (**Figures 2D and S2F**). No overall difference in weight gain or organ mass was observed in genetic control mice lacking caspase-8 only in myeloid cells (*Casp8^LysMcre^*) that display inefficient deletion on a RIPK3 sufficient background (**Figures S2G and S2H**)^31^.

**Figure 2.**
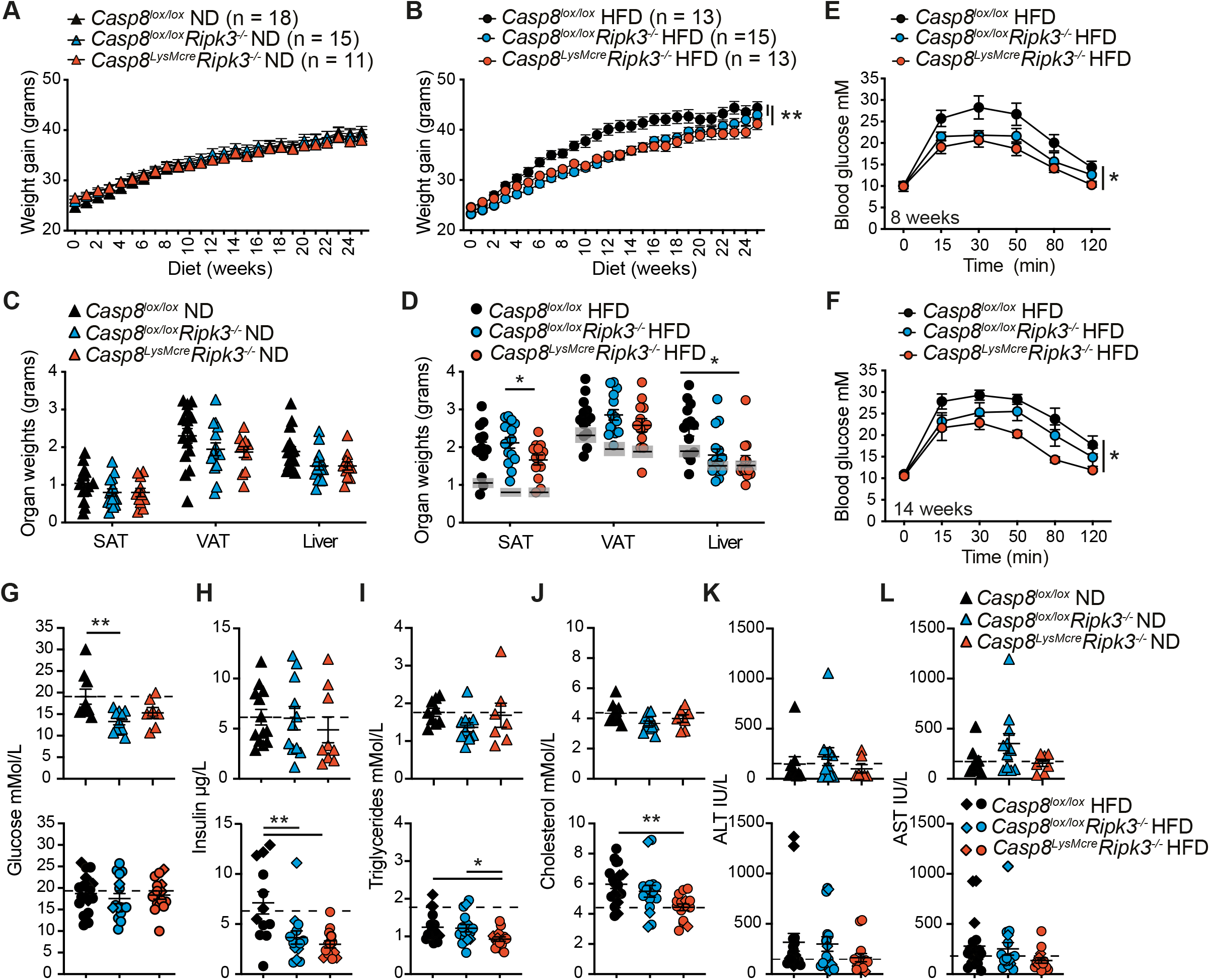
Myeloid specific RIPK3-caspase-8 signaling drives HFD-induced metabolic dysfunction. **(A, B)** Body weights were measured weekly in *Casp8^lox/lox^, Casp8^lox/lox^Ripk3^-/-^*, and *Casp8^LysMcre^Ripk3^-/-^* mice fed a **(A)** normal chow diet (ND) or **(B)** high-fat diet (HFD) for ~ 25 weeks. (ND n ≥ 11 mice per group and HFD n ≥ 13 mice per group pooled from 3 independent experiments). **(C, D)** End stage organ weights at in **(C)** ND- and **(D)** HFD-fed *Casp8^lox/lox^, Casp8^lox/lox^Ripk3^-/-^*, and *Casp8^LysMcre^Ripk3^-/-^* mice. (n ≥ 11 mice per group pooled from 3 independent experiments). Grey box in **(D)** shows the mean ± SEM from **(C)** for comparisons. **(E, F)** Glucose tolerance was measured in HFD challenged *Casp8^lox/lox^*, *Casp8^lox/lox^Ripk3^-/-^*, and *Casp8^LysMcre^ Ripk3^-/-^* mice by intraperitoneal glucose tolerance tests (IP-GTT; 1.5 g/kg) at 8 and >14 weeks. (n = 5-6 mice per group). Data are representative of one of 2-3 experiments. **(G-L)** Fasting serum **(G)** glucose, **(H)** insulin, **(I)** triglycerides, **(J)** cholesterol, **(K)** ALT and **(L)** AST levels in *Casp8^lox/lox^, Casp8^lox/lox^Ripk3^-/-^*, and *Casp8^LysMcre^Ripk3^-/-^* mice after 16-18 (Diamonds) or 25 weeks (Triangles/circles). (n ≥ 9 mice per group pooled from at least 3 experiments). Dotted line shows mean values of control *Casp8^lox/lox^* ND-fed mice extrapolated from top panel. Data are the mean ± SEM. *p < 0.05; **p < 0.01; ***p < 0.001; ****p < 0.0001. One-way ANOVA of AUC (A, B, E, F) and One-way ANOVA (C, D, G-L). See related **Figure S2.**

Examination of glycemic control revealed that ND- and HFD-fed *Casp8^lox/lox^Ripk3^-/-^* and *Casp8^LysMcre^Ripk3^-/-^* mice had comparable fasting blood glucose levels when compared to controls (**Figures 2E, 2F and S2I-S2K**). ND-fed animals exhibited no significant difference in glucose clearance during an intraperitoneal-glucose tolerance test (IP-GTT) after 8 or 16 weeks of diet (**Figures S2I and S2J**). In contrast, HFD-fed *Casp8^LysMcre^Ripk3^-/-^* mice demonstrated superior glucose clearance over-time, compared to control and RIPK3-deficient mice (**Figures 2E, 2F and S2K**), but were equally resistant to insulin after ~23 weeks of HFD (**Figure S2L**). Although, at endpoint, HFD-fed *Casp8^lox/lox^Ripk3^-/-^* and *Casp8^LysMcre^Ripk3^-/-^* mice exhibited lower starved insulin levels (**Figures 2G and 2H**), which may be indicative of better insulin sensitivity. Strikingly, examination of other systemic metabolic disease markers revealed HFD-fed *Casp8^LysMcre^Ripk3^-/-^* mice exhibited reduced signs of dyslipidemia, with significantly lower serum triglyceride and cholesterol levels upon HFD feeding (**Figures 2I and 2J**). In contrast, serum ALT and AST levels (as markers of liver damage) were not significantly attenuated in the HFD-fed *Caspase-8^LysMcre^Ripk3^-/-^* cohort (**Figures 2K and 2L**). Of note, HFD-fed RIPK3-deficient mice partially phenocopied *Caspase-8^LysMcre^Ripk3^-/-^* mice with trends toward lower insulin, triglyceride, and cholesterol (**Figures 2H–2J**). Together, these data indicate that caspase-8 contributes to obesity-induced metabolic dysfunction.

### Loss of RIPK3 and caspase-8 activity in myeloid cells reduces local inflammation and steatosis

The accumulation of monocyte-derived CD11c^+^F4/80^+^ macrophages in adipose tissue and formation of crown-like structures around dying adipocytes is believed to drive the chronic low-level inflammation that leads to metabolic dysfunction and insulin resistance^56^. We therefore examined whether RIPK3 and caspase-8 contribute to pathological inflammatory changes in HFD-induced obesity. In line with comparable VAT weights between HFD-fed groups (**Figures 2D and S2F**), the mean adipocyte size from HFD-fed *Casp8^LysMcre^Ripk3^-/-^* mice was similar to control mice (**Figures 3A, 3B and S2M**). Yet, intriguingly, fewer F4/80^+^ crown-like structures (**Fig. 3A**), less monocyte/macrophage and neutrophil infiltrate (**Figures 3C and S2N**), and reduced IL-1 β and TNF secretion (to LPS treatment) (**Figure 3D**) were evident in the VAT of HFD-fed *Casp8^LysMcre^Ripk3^-/-^* mice and, to a lesser degree *Casp8^lox/lox^Ripk3^-/-^* mice. Histopathologic analysis of the livers also revealed that myeloid-specific caspase-8 deficiency with RIPK3-loss ameliorated hepatic steatosis, ballooning and fibrosis in HFD-fed (partial effect in ND-fed) mice (**Figures 3E and 3F**), whilst flow cytometric analysis suggested reduced recruitment of inflammatory monocytes and neutrophils in *Casp8^lox/lox^Ripk3^-/-^* and *Casp8^LysMcre^Ripk3^-/-^* HFD-fed mice (**Figures 3G and 3H**). Collectively, these results together with our *in vitro* findings (**Figures 1E, 1G and S2I-S2K**), suggest that myeloid cell caspase-8 activity can drive adipose tissue inflammation and the progression to NASH.

**Figure 3.**
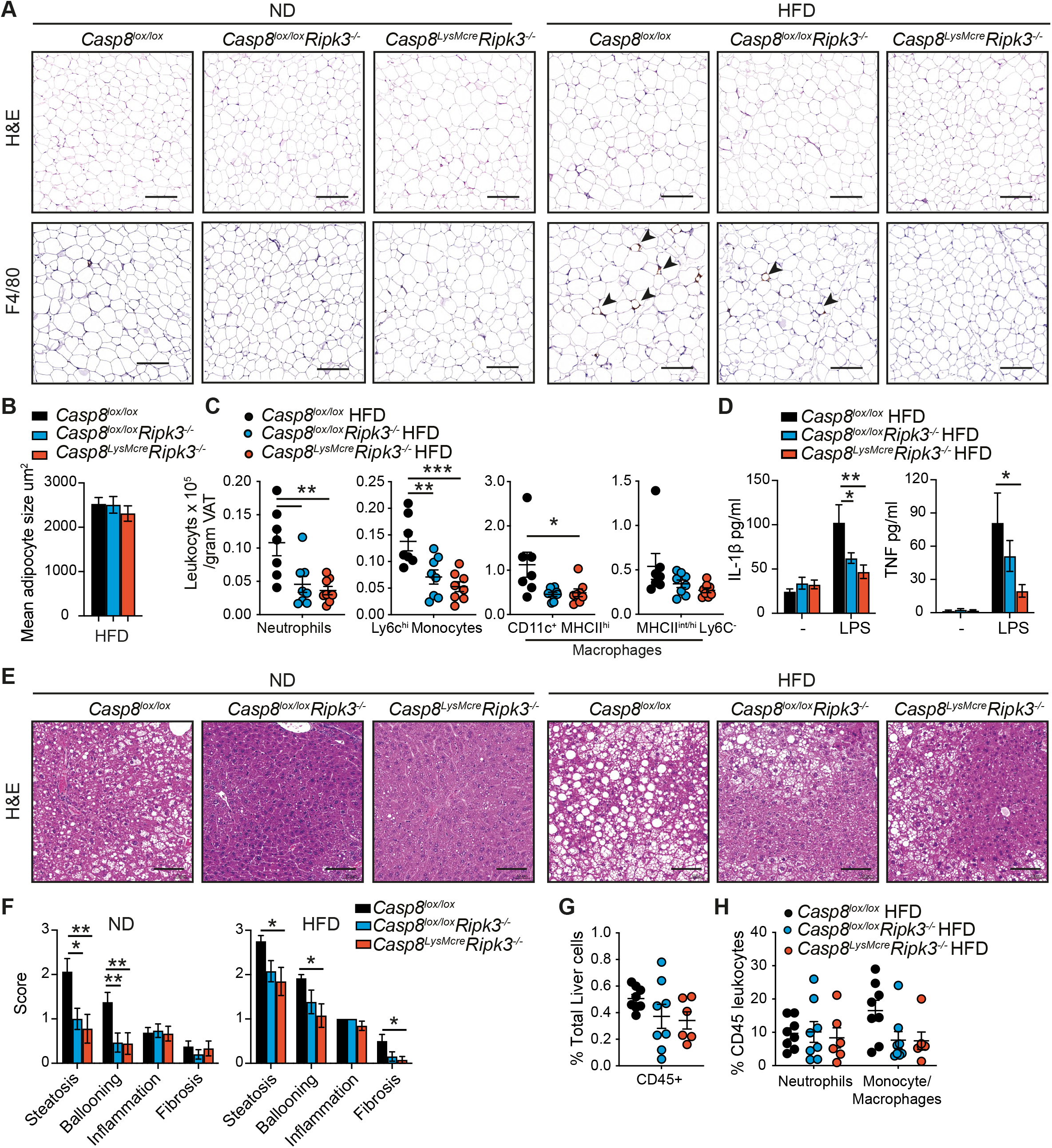
Myeloid specific RIPK3-caspase-8 signaling promotes HFD-induced inflammation and NAFLD. **(A)** Representative H&E stained and F4/80 immuno-stained VAT tissue sections from ND- and HFD-fed *Casp8^lox/lox^*, *Casp8^lox/lox^Ripk3^-/-^*, and *Casp8^LysMcre^Ripk3^-/-^* mice at 25 weeks. Arrow points to crownlike structures. Scale bar is 0.2 mm **(B)** Mean adipocyte size in HFD-fed *Casp8^lox/lox^, Casp8^lox/lox^Ripk3^-/-^*, and *Casp8^LyMcre^Ripk3^-/-^* VAT was quantified using an automated algorithm. (n ≥ 11 mice per group pooled from 3 independent experiments). **(C)** Numbers of neutrophils, inflammatory monocytes and macrophages were quantified in the VAT tissue of HFD-fed challenged *Casp8^lox/lox^*, *Casp8^lox/lox^Ripk3^-/-^*, and *Casp8^LysMcre^Ripk3^-/-^* mice by flow cytometric analysis. (n = 7-8 mice per group pooled from 2 experiments). **(D)** VAT from HFD-fed *Casp8^lox/lox^, Casp8^lox/lox^Ripk3^-/-^*, and *CRsp8^LysMcre^Ripk3^-/-^* mice was cultured *ex vivo* with and without LPS (50 ng/ml) overnight and IL-1β and TNF measured in the supernatants by ELISA. (n ≥ 12 mice per group pooled from 3 experiments). **(E, F)** Representative **(E)** H&E stained liver sections (scale bar is 0.1 mm) and **(F)** histopathological evaluation of disease in *Casp8^lox/lox^*, *Casp8^lox/lox^Ripk3^-/-^*, and *Casp8^LysMcre^Ripk3^-/-^* ND-fed and HFD-fed mice after 25 weeks of challenge. (n ≥ 9 mice per group pooled from 3 experiments). **(G, H)** Flow cytometric analysis of monocyte/macrophage and neutrophils in the livers of *Casp8^lox/lox^*, *Casp8^lox/lox^Ripk3^-/-^*, and *Casp8^LysMcre^ Ripk3^-/-^* HFD mice. (n = 6-8 mice per group pooled from 2 experiments). Data are the mean ± SEM. *p < 0.05; **p < 0.01; ***p < 0.001; ****p < 0.0001. One-way ANOVA. See related **Figure S2.**

### MLKL drives obesity-induced metabolic dysfunction and insulin resistance

We next examined the effect of RIPK3 substrate MLKL on obesity and NAFLD. ND-fed MLKL-deficient mice displayed steady weight gain, although weights peaked lower than WT control mice (**Figures 4A and S3A**) and was associated with a reduction in both SAT and VAT weights (**Figures 4B and S3C, S3E**). Impressively, HFD-fed *Mlkl^-/-^* mice exhibited markedly diminished weight gain over-time, compared to controls (**Figures 4C and S3B**), which correlated with reduced SAT, VAT and liver tissue expansion (**Figures 4D and S3D, S3E**), and not with altered food intake or fecal output (**Figure S3F**). Upon testing glucose and insulin resistance, *Mlkl^-/-^* mice demonstrated less metabolic dysfunction on a HFD diet, and to a lesser extent on a ND (**Figures 4E–4I and S3G**). Analysis of metabolic and damage markers in ND-fed mice revealed that fasting serum glucose, insulin, ALT, AST, triglyceride, cholesterol and non-esterified fatty acids (NEFA) levels were comparatively normal in *Mlkl^-/-^* mice (**Figures 4J–4P**, **top panel**). However, upon HFD feeding, *Mlkl^-/-^* mice exhibited significantly reduced serum insulin, cholesterol and ALT levels, compared to HFD-fed WT controls (**Figures 4J–4P**, **bottom panel**); supporting that MLKL drives obesity-induced metabolic dysfunction.

**Figure 4.**
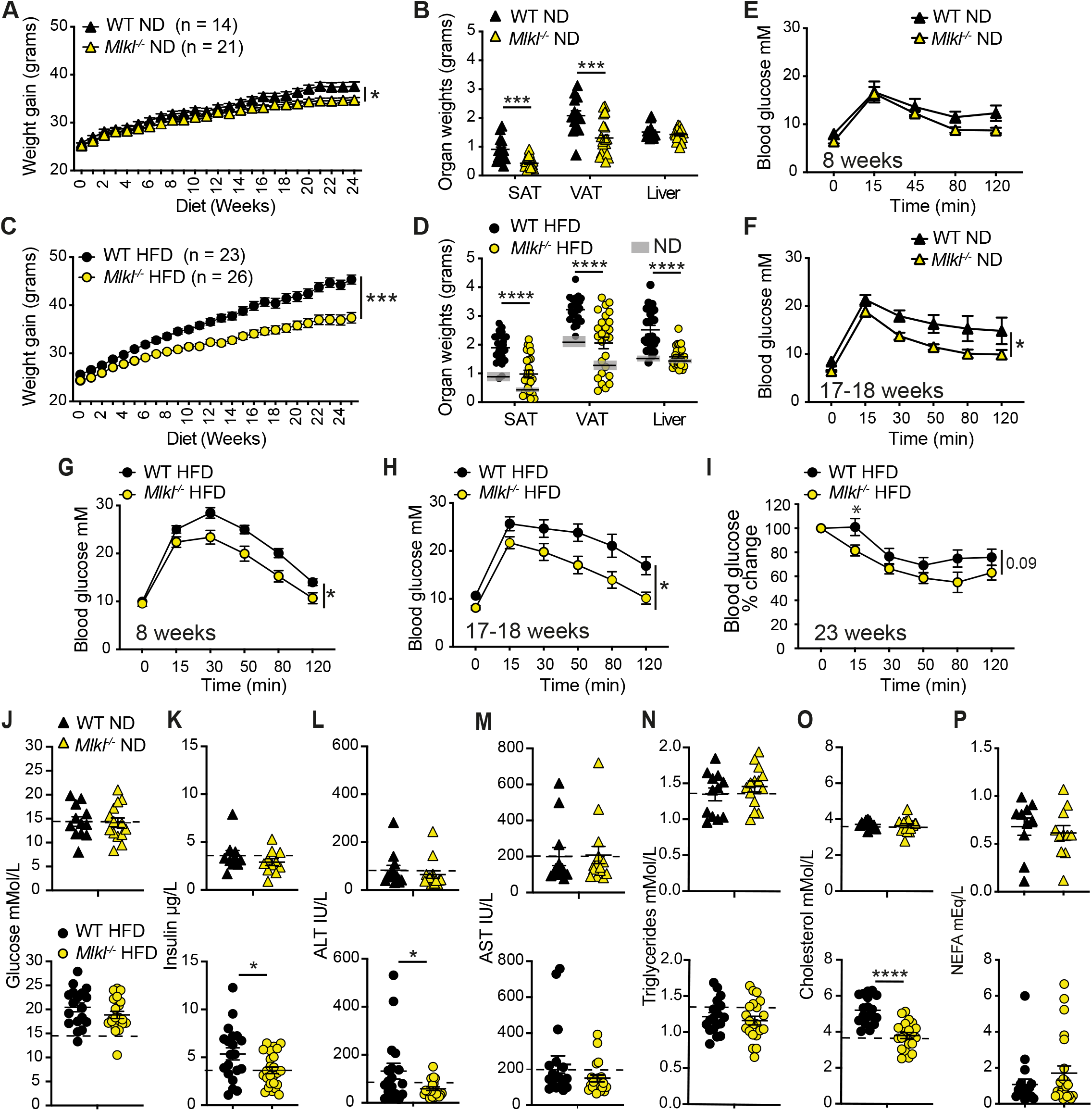
MLKL signaling exacerbates HFD-induced obesity and metabolic dysfunction. **(A, C)** WT and *Mlkl^-/-^* mice were fed a **(A)** ND- and **(C)** HFD for ~ 24-25 weeks and body weights measured on a weekly basis. (ND n ≥ 14 mice/group and HFD n ≥ 23 mice/group pooled from 3 independent experiments). **(B, D)** End stage organ weights from **(B)** ND- and **(D)** HFD-fed WT and *Mlkl^-/-^* mice. (n ≥ 14 mice/group pooled from 3 experiments). Grey box in **(D)** represents the mean ± SEM from **(B)** for comparison. **(E-H)** Glucose tolerance was assessed via an IP-GTT (1.5 g/kg) at 8 and 17-18 weeks in **(E, F)** ND- (n = 5-6 mice per group) and **(G, H)** HFD-fed (n = 8-9 mice per group) WT and *Mlkl^-/-^* mice. Data are representative of one of 2-3 experiments. **(I)** Insulin resistance was assessed in HFD-fed WT and *Mlkl^-/-^* mice (~ 23 weeks) during IP-ITT (0.75 U/kg). (n = 8-9 mice/group from one representative experiment of 2 experiments). **(J-P)** Fasting serum **(J)** glucose, **(K)** insulin, **(L)** ALT, **(M)** AST, **(N)** triglyceride, **(O)** cholesterol and **(P)** NEFA after 24-25 weeks of ND (n ≥ 10 mice per group, top panel) or HFD (n ≥ 18 mice per group, bottom panel). Data are from 3 pooled experiments. Data are the mean ± SEM. *p < 0.05; **p < 0.01; ***p < 0.001; ****p < 0.0001. Student’s T-test of AUC (A, C, E-H) and Student’s T-test (B, D, J-P). See related **Figure S3.**

### MLKL regulates adiposity and the development of NAFLD

We next examined VAT and liver pathologies to discern how MLKL drives obesity and liver disease. Correlating with reduced VAT weights (**Figures 4B and 4D**), ND and HFD-fed *Mlkl^-/-^* adipocytes were smaller on average (**Figures 5A and 5B**), suggesting that MLKL regulates adiposity. Surprisingly, analysis of inflammation in VAT tissue from HFD-fed mice revealed only a nonsignificant trend towards reduced F4/80^+^ crown-like macrophage structures and inflammatory cell infiltrate in MLKL-deficient mice, compared to controls (**Figures 5A and 5C**). Correspondingly, LPS-induced IL-1β and TNF cytokine secretion in VAT was only modestly impaired (**Figure 5D**). Analysis of liver pathology also revealed that MLKL-deficient mice were markedly protected from steatosis and ballooning from HFD-feeding, with a similar trend in aging ND-fed mice (**Figures 5E and 5F**). *Mlkl^-/-^* livers also exhibited a select reduction in the proportion of monocyte/macrophages within the liver tissue infiltrate, although this reduction was associated with reduced expression of *Tnfa* and not inflammasome-associated genes, (**Figures 5G and 5H**). Therefore, MLKL activity drives obesity and NAFLD in aged and HFD-fed mice but does not contribute substantially to inflammation.

**Figure 5.**
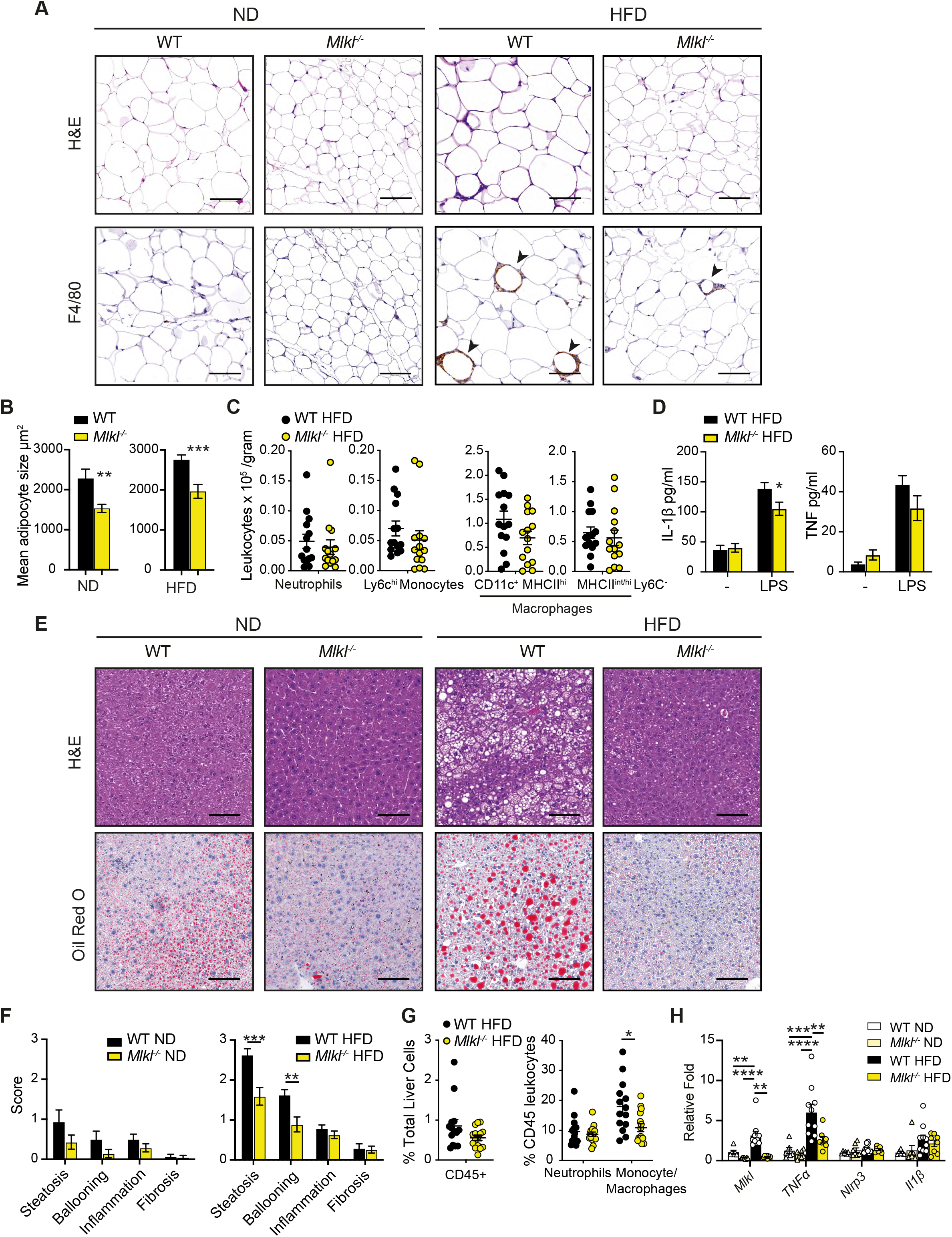
MLKL deficiency reduces adiposity and fatty liver disease in response to HFD challenge. **(A, B)** Representative **(A)** H&E stained and F4/80 immuno-stained VAT tissue sections from WT and *Mlkl^-/-^* mice fed a ND or HFD for ~25 weeks, and **(B)** automated quantification of mean adipocyte size in VAT. Arrows point to crown-like structures. Scale bars are 0.1 mm. (ND n ≥ 13 mice per group and HFD n ≥ 23 mice per group from 3 pooled experiments). **(C)** VAT from HFD-fed WT and *Mlkl^-/-^* mice was harvested at ~25 weeks and numbers of neutrophils, inflammatory monocytes and macrophages quantified by flow cytometric analysis. (n ≥ 14 mice per group pooled from 2 experiments). **(D)** VAT from HFD-fed WT and *Mlkl^-/-^* mice was harvested at ~25 weeks and cultured *ex vivo* with and without LPS (50 ng/ml) overnight and IL-1β and TNF measured in the supernatants by ELISA. (n ≥ 19 mice per group from 3 pooled experiments). **(E, F)** Representative **(E)** H&E stained and Oil red O stained liver sections (scale bars are 0.1 mm) and **(F)** histopathological evaluation of disease in WT and *Mlkl^-/-^* mice after 25 weeks of ND or HFD feeding. (n ≥ 14 ND-fed mice and n ≥ 23 HFD-fed mice pooled from 3 cohorts). **(G)** Flow cytometric analysis of the proportion of CD45^+^ leukocytes in the in the livers of WT and *Mlkl^-/-^* HFD mice that are monocyte/macrophage and neutrophils. (n ≥ 14 mice per group pooled from 2 experiments, mean ± SEM. **(H)** qRT-PCR gene expression in ND and HFD liver tissue after 23-25 weeks of diet. (n ≥ 6 mice per group pooled from 3 experiments). Data are the mean ± SEM. *p < 0.05; **p < 0.01; ***p < 0.001; ****p < 0.0001. Student’s T-test. See related **Figure S3.**

To evaluate whether hematopoietic MLKL expression contributes to obesity, we examined C57BL/6 mice reconstituted with WT control or *Mlkl^-/-^* bone marrow in our dietary model. Both weekly weighing and measurement of fat mass using an EcoMRI instrument highlighted that WT and *Mlkl^-/-^* bone marrow chimeras responded equivalently to a ND or HFD (**Figures S3H and S3I**). Furthermore, in contrast to global MLKL-deficient mice, no differences in end-stage SAT, VAT or liver weights were observed in chimeric animals (**Figure S3J**) nor were there signs of improved glucose metabolism (**Figure S3K**). These results, combined with the reduced adipocyte hypertrophy and liver damage observed in aged or HFD-fed MLKL-deficient mice, suggests that MLKL activity alters tissue homeostasis to cause obesity-induced metabolic disease.

### MLKL signaling induces lipid metabolic gene signatures in the liver of ageing and obese mice

In view of the protection from NAFLD and metabolic dysfunction observed in MLKL-deficient mice, we next examined global hepatic responses in aged ND-fed or HFD-fed mice. 3’ mRNA sequencing of WT and *Mlkl^-/-^* livers revealed that there were significant transcriptomic changes in differentially expressed genes (DEGs) between genotypes and diets (**Figures 6A and 6B**). In WT HFD-fed mice, more than 1700 genes (p<0.05) were up- and down-regulated, while *Mlkl^-/-^* HFD mice exhibited 1111 upregulated and 948 downregulated genes, compared to ND mice (p<0.05). As expected, following HFD-feeding, gene ontology (GO) analyses revealed upregulation of metabolic and inflammatory processes in WT livers, such as cholesterol biosynthesis, acyl-CoA metabolic process, Peroxisome proliferator-activated receptor (PPAR) signaling pathway and inflammatory response (**Figure S4A**).

**Figure 6.**
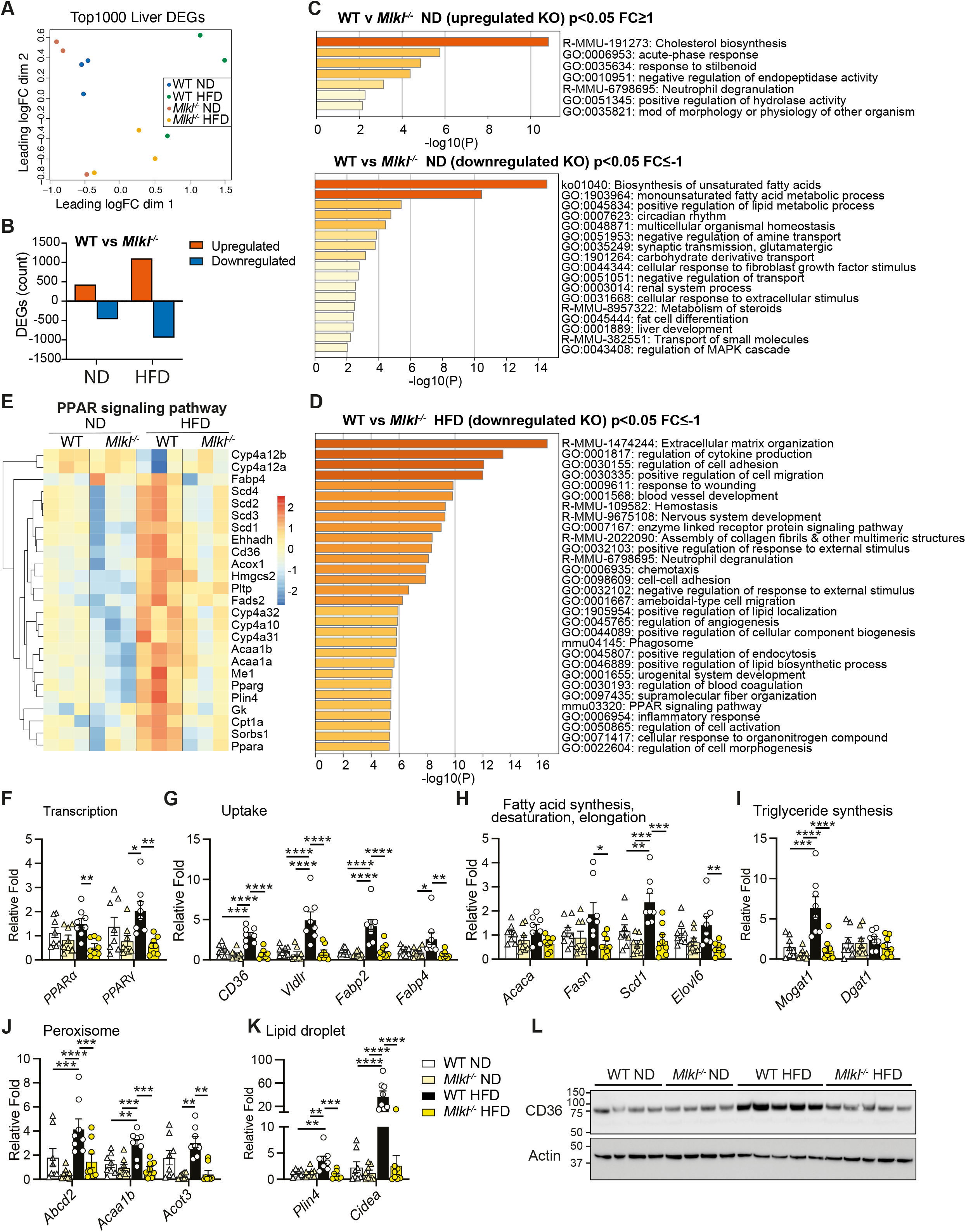
MLKL regulates genes involved in lipid metabolism in the liver. **(A-E)** Liver RNA extracted from WT and *Mlkl^-/-^* mice on a ND or HFD (n = 3 mice per group) was subjected to 3’ mRNA sequencing. **(A)** Multidimensional scaling (MDS) plot and **(B)** Number of differentially expressed genes (DEGs) upregulated and downregulated in WT v *Mlkl^-/-^* ND and HFD livers. P ≤ 0.05 and cut-off values logFC ≥ 1 or logFC ≤ −1. **(C)** Gene Ontology (GO) pathways of significant DEGs upregulated and downregulated in *Mlkl^-/-^* ND livers compared to WT ND livers. p ≤ 0.05 and cut-off values logFC ≥ 1 or logFC ≤ −1. **(D)** Top 30 GO pathways of significant DEGs downregulated in *Mlkl^-/-^* HFD livers compared to WT HFD livers. p ≤ 0.05 and cut-off values logFC ≤ −1. **(E)** Heatmap of significant DEGs of GO term PPAR signaling pathway (downregulated in *Mlkl^-/-^* HFD livers compared to WT HFD livers). p ≤ 0.05 and cut-off values logFC ≥ 1 or logFC ≤ −1. **(F-K)** qRT-PCR analysis of liver mRNA from WT and *Mlkl^-/-^* mice fed a ND or HFD. (n = 6-8 mice per group from 3 experiments). **(L)** Liver lysates from ND- and HFD-fed WT and *Mlkl^-/-^* mice were analyzed by immunoblot for the indicated antibodies. Each lane represents an individual mouse (n = 4-5 mice per group). Data are the mean ± SEM. *p < 0.05; **p < 0.01; ***p < 0.001; ****p < 0.0001. Student’s T-test. See related **Figure S4 and S5**.

Likewise, livers from HFD-fed versus ND-fed *Mlkl^-/-^* mice showed upregulation of lipid metabolism-associated terms (*e.g*., fatty acid metabolic process, long-chain fatty acid metabolic process) (**Figure S4B**). Importantly, direct comparisons between WT and *Mlkl^-/-^* livers revealed that MLKL deficiency drives significant changes in DEGs in HFD mice, and modestly in ND (**Figure 6B**).

Gene set enrichment analysis (GSEA) and GO analysis revealed that, in the absence of MLKL, genes associated with cholesterol biosynthesis were increased in the ageing liver (**Figures 6C and S4D**). Conversely, GO analyses revealed that ND-fed *Mlkl^-/-^* livers exhibited downregulated gene sets for various lipid-associated and metabolic processes, including biosynthesis of unsaturated fatty acids and positive regulation of lipid metabolic process (**Figure 6C**). GSEA further supported this trend, showing downregulation of genes associated with oxidative phosphorylation, fatty acid metabolism, peroxisomes and adipogenesis (**Figure S4D**). Importantly, HFD-fed *Mlkl^-/-^* mice failed to upregulate these gene signatures, with the exception of oxidative phosphorylation that was upregulated compared to HFD-fed WT livers and characterized by increased electron transport chain gene expression (**Figure S4C, S4E and S5A**). GO analysis further complemented this pattern with downregulation of gene sets involved in several lipid and membrane regulatory/signaling processes in HFD-fed *Mlkl^-/-^* livers, such as regulation of cytokine production, positive regulation of lipid localization, positive regulation of lipid biosynthetic process and PPAR signaling pathway (**Figure 6D and S5B-S5D**). Notably, many genes downregulated in HFD MLKL-deficient livers were extracellular or intracellular sensors/receptors that coordinate cell cytokine/signaling responses (**Figure S5B**).

Comparisons of downregulated lipid/membrane-related GO terms in HFD-fed *Mlkl^-/-^* livers revealed an overlap in genes regulated by nuclear PPAR signaling (**Figures 6D, 6E and Figure S5B-S5D**). Correspondingly, qRT-PCR analysis validated that several PPAR-related genes that were differential in GO analyses (**Figures 6E and S5B-S5D**) and/or are associated with key lipogenic processes were reduced in *Mlkl^-/-^* livers with HFD feeding (and trended down in ND-fed animals), including the transcription factors *Pparα/γ* themselves, as well as genes associated with fatty acid uptake (*Cd36, Fabp4, Vldlr*), fatty acid synthesis (*Acaca, Fasn*), elongation (*Elovl6*) and desaturation (*Scd1*), triglyceride synthesis (*Mogat1, Dgat1*), peroxisome function (*Acot3, Abcd2, Acaa1b*), lipid droplet storage (*Plin4*) and death/lipolysis (*Cidea*) (**Figures 6F–6K**). As CD36 is a key membrane fatty acid translocase that triggers a PPAR-regulated positive-feedback loop and several other lipid homeostatic processes^57–59^, we assessed CD36 levels by immunoblot and uncovered that HFD-fed *Mlkl^-/-^* livers failed to upregulate CD36 protein levels (**Fig. 6L**). Collectively, these findings suggest that MLKL signaling regulates molecules involved in lipid uptake/transport, synthesis and signaling.

### MLKL regulates the synthesis of monounsaturated and polyunsaturated diglycerides and triglycerides in the liver

The active N-terminal four-helical bundle of MLKL dominantly binds negatively charged phosphatidyl-inositol-phosphates (PIPs), particularly PI(4,5)P2, in the plasma membrane to facilitate necroptotic death^19,60,61^. Beyond this, necroptosis activation and signaling may be tightly regulated by lipid species^62^, as saturated very long chain fatty acids (VLCFAs) and acylation of phospho-MLKL/MLKL is required to promote endocytic trafficking of MLKL to the membrane^63,64^. Based on the perturbed lipid metabolic/membrane gene signatures in the livers of MLKL-deficient mice, we sought to understand how MLKL signaling impacts the abundance of individual lipid species in the serum, VAT and liver of ageing and HFD-diet fed mice using targeted lipidomic analysis. Lipid metabolic clustering was primarily associated with diet but some divergence of serum and liver profiles between genotypes was observed on HFD, with a modest shift in hepatic lipids also seen with ND-feeding (**Figure S6A**). In contrast, only a minor shift in lipid profiles was observed in the VAT of HFD-fed WT and MLKL-deficient mice (**Figure S6A**).

Analysis of the relative abundance of individual lipid species and/or classes in the serum, liver and VAT of ageing ND-fed WT and *Mlkl^-/-^* mice revealed no major changes (**Figures 7A, 7B** and **S6A)**. Despite no gross perturbations in lipid signature in the VAT of ND- or HFD-fed WT and *Mlkl^-/-^* mice, specific analysis of obesity-associated ceramide (CER) species, revealed a downward trend in most species in HFD-fed *Mlkl^-/-^* VAT, including toxic long chain fatty acid species (**Figure S6B**)^65^. Analysis of the relative abundance of lipid classes in the serum of HFD-fed WT and *Mlkl^-/-^* mice also revealed no overall changes in total Diglycerides (DG) and Triglycerides (TG) in HFD-fed WT and *Mlkl^-/-^* serum (**Figure 7A**). However, levels of select monounsaturated fatty acid (MUFA) and polyunsaturated fatty acid (PUFA) TG species were significantly elevated, and a trend towards increased saturated TG was also evident (**Figures 7C and S6C**), perhaps indicative of active secretion and/or reduced uptake by metabolic tissues. Intriguingly, membrane lipids (*e.g*., sphingomyelin (SM), phosphatidylcholine (PC) and phosphatidylinositol (PI)), particularly those comprised of long to very long acyl chains with at least one double bond, were reduced in HFD-fed *Mlkl^-/-^* serum (**Figures 7A and 7C**).

**Figure 7.**
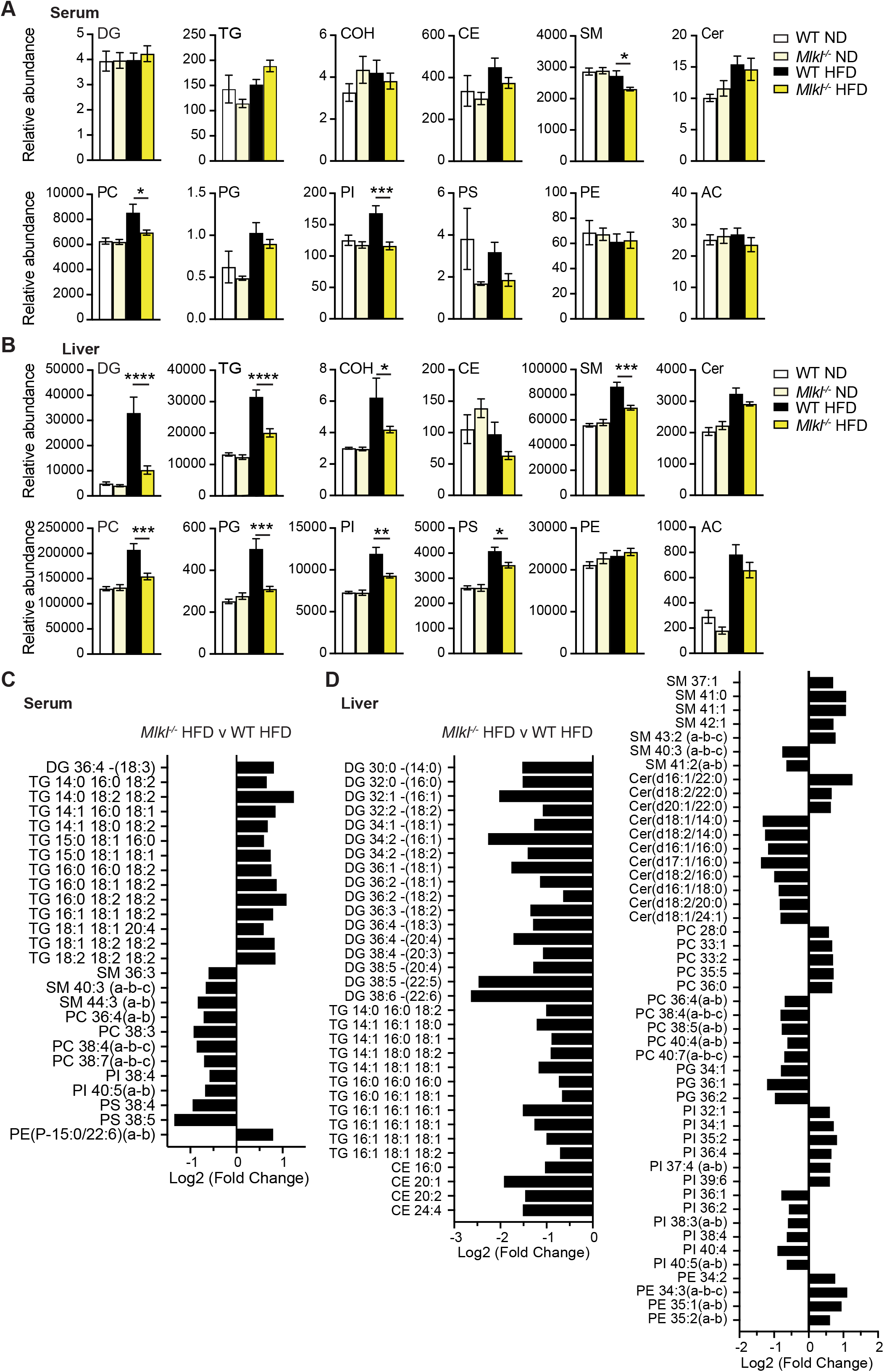
MLKL deficiency alters lipid species in the serum and liver upon HFD feeding. Wildtype (WT) and *Mlkl^-/-^* were fed a normal chow diet (ND) or high fat diet (HFD) for 25 weeks. Total lipid was extracted from the serum, liver and VAT and lipid species analyzed by LCMS. **(A, B)** Relative abundance of total lipid classes in the **(A)** serum and **(B)** liver of mice. Data are normalised for the median lipid content per mouse and, where relevant, for tissue mass. (n = 8-12 mice per group pooled from 3 experiments). Statistical analyses shown were calculated using the median lipid/tissue weight normalised data after (log10) transformation. **(C, E)** Fold change (log2) of individual lipid species in *Mlkl^-/-^* HFD-versus WT HFD-fed **(C)** serum and **(D)** liver. Data shown are median normalised log2 transformed data adjusted for a false discovery rate (unpaired Student’s T-test p < 0.05). Key: Diglycerides (DG) Triglycerides (TG), Cholesterol esters (CE), Ceramide (Cer), Sphingomyelin (SM), Acylcarnitine (Ac) Phosphatidylcholine (PC), Phosphatidylinositol (PI), Phosphatidylserine (PS), Phosphatidylglycerol (PG), Phosphatidylethanolamine (PE), and Cholesterol (COH). Data are the mean ± SEM. *p < 0.05; **p < 0.01; ***p < 0.001; ****p < 0.0001. One-way ANOVA followed by Tukey’s multiple comparisons test was applied (A, B) and Student’s T-test (C, D). See related **Figure S6**.

Liver lipid profiling revealed that levels of DG were attenuated in *Mlkl^-/-^* livers upon HFD feeding (**Figures 7B, 7D and S6D**). Correspondingly, pools of TG species and phospholipids PC, PI, PS, phosphatidylglycerol (PG), which are dependent on DG for their synthesis, were significantly reduced in HFD-fed *Mlkl^-/-^* livers, as were SM species (**Figures 7B and D**). Closer examination of the DG and TG composition revealed a reduction in the abundance of saturated TG comprising three palmitic acids (C16:0) in *Mlkl^-/-^* livers, and a diminished abundance of MUFA or PUFA DG, TG and phospholipid species containing at least one long chain (oleic acid C18:1, arachidonic acid C20:4) or very long chain (nervonic acid C24:1, docosahexaenoic acid C22:6) fatty acid (**Figures 7D and S6D**). This pattern suggests that MLKL-deficient mice may be defective in the uptake and/or synthesis/remodelling of unsaturated long and very long acyl chain fatty acids. Moreover, these results correlate strongly with our RNA-seq data set, and further substantiate that MLKL regulates lipid homeostasis to drive obesity and the development of fatty liver disease.

## DISCUSSION

In this study, we genetically examined the contribution of extrinsic cell death signaling to high-fat diet (HFD)-induced obesity and associated metabolic dysfunction. Our findings show that extrinsic caspase-8 activity in macrophages largely induces chronic metabolic inflammation by regulating the transcriptional responses needed for efficient NLRP3 inflammasome activity. In comparison, MLKL appears to drive obesity with ageing to promote NAFLD development in a manner independent of canonical RIPK3 signaling. Remarkably, the protection from NAFLD we observed in MLKL-deficient mice appeared to be conferred by reduced signals for lipid uptake, *de novo* lipogenesis and triglyceride synthesis and culminated in a deficit in long chain and very long chain unsaturated lipid species.

The NLRP3 inflammasome has emerged as an attractive therapeutic target to limit pathogenic IL-1 β activity in obesity-associated metabolic disorders^7^. A large body of evidence places caspase-8 or MLKL signaling upstream of NLRP3 inflammasome activity in a range of disease settings^44,54,66–69^ yet, their contribution to NLRP3 inflammasome activity in metabolic disease is less clear^20^. Our findings reveal that while myeloid-specific caspase-8 and global RIPK3 loss mostly phenocopies the protection from adipose tissue inflammation, metabolic dysfunction and steatohepatitis observed in NLRP3 inflammasome-deficient mice^8–10^, it failed to prevent adiposity and end-stage insulin resistance. Accordingly, we uncovered that RIPK3-caspase-8 signaling is not obligatory for NLRP3 inflammasome activation in LPS-treated macrophages exposed to palmitate, but instead caspase-8 regulates inflammatory gene transcription, direct IL-1 β proteolysis, and to a lesser extent cell death in LPS and palmitate treated macrophages. Regarding the latter, it is possible that saturated fatty acid crystallization in the autophagosome^70^ ultimately leads to autophagolysosomal rupture and cellular demise, as observed for other crystalline NLRP3-activating stimuli^71^. Overall, our results suggest that targeting apoptotic caspase-8 activity may potentially limit myeloid cell-driven tissue inflammation and subsequently dampen NAFLD progression. However, the utility of this approach may be limited given the poor clinical performance of the pan-caspase inhibitor emricasan in NAFLD/NASH^15^.

While the necroptotic kinases, RIPK1 and RIPK3, have been studied in liver damage models^72^, it remains controversial as to whether antagonizing these kinases and necroptotic signaling is a valid therapeutic option in obesity-driven NAFLD/NASH, as disease outcomes differ based on the dietary intervention and genetic models used^14,20,22–26,73,74^ For example, RIPK3 deficiency partially protects mice from NASH development on a MCD-diet and a choline-deficient diet, but not a HFD^14,23,25,26^. Intriguingly, recent reports have argued that necroptosis in hepatocytes is limited *via* the epigenetic suppression of RIPK3^75,76^. Yet, the elevated RIPK1, RIPK3 and MLKL expression observed in progressive liver injury and severe NALFD/NASH provides evidence that hepatocytes may overcome this defect in certain contexts^14,26,75,77^, or that, alternatively, RIPK3-MLKL signaling may be active in disease-promoting parenchymal cholangiocytes and/or non-parenchymal cells^14^. Importantly, our work suggests that both RIPK3 and MLKL do indeed contribute to NAFLD progression but that MLKL has divergent activities in obesity and metabolic dysfunction. Specifically, MLKL deletion reduced adipose tissue hypertrophy, and afforded greater protection from metabolic dysfunction, insulin resistance and NAFLD, when compared to RIPK3-deficient mice. Unexpectedly, MLKL loss did not appear to markedly attenuate inflammatory changes (*i.e*., cytokine secretion and inflammatory cell influx) in adipose and liver tissues to HFD feeding, contrasting with other reports^27,29^. The reason for discrepancies between studies is unclear but is likely to be influenced by the dietary model adopted and duration of challenge, genetic background and use of littermates, and environmental variations (*e.g*., microbiome, housing).

MLKL appears to act independent of canonical RIPK3 signaling in obesity and metabolic dysfunction. How MLKL protects from obesity is unclear but one recent study proposed that MLKL, and not RIPK3, drives white adipose tissue differentiation^78^. MLKL loss also distinctly caused a select reduction in circulating cholesterol, akin to a recent report in a model of atherosclerosis^79^. However, while we observed this phenomenon was associated with reduced lipid accumulation in tissues, Rasheed *et al*. observed that MLKL promotes the retention of lipid in macrophages within the atherosclerotic plaque^79^, suggesting differences in MLKL functions between cell types. MLKL has also been acknowledged to have noncanonical actions in the liver, including the inhibition of autophagic flux, insulin signaling and mitochondrial biogenesis, as well as in increasing *de novo* lipogenesis^21,27–29^. Our RNA-seq analysis revealed a dominant role for MLKL in perturbing lipid homeostasis in the liver, although, we do not discount the possibility that these other processes contribute to disease. In particular, we observed downregulation of a number of key PPARα/γ-induced genes that regulate lipid uptake, transport, synthesis and storage in the HFD-fed *Mlkl^-/-^* liver. It remains unclear how MLKL regulates the transcription of genes involved in lipid metabolism, although, it is likely a consequence of MLKL’s propensity to target lipid-rich endosomes, autophagolysosomes and plasma membranes^27,64,79,80^. We also observed that, in the absence of MLKL, there was reduced expression of the fatty acid uptake receptor CD36 in the liver, which can signal *via* PPARs, and plays a key in promoting *de novo* lipogenesis and limiting β-oxidation and autophagy^81,82^. As normal CD36 expression has been reported in atherosclerotic macrophage foam cells lacking MLKL^79^, our work again highlights that MLKL may differentially regulate lipid regulatory molecules and thus affect lipid handling and storage differently depending on the pathogenic cell type.

Closely aligning with our transcriptomic data, targeted lipidomics of tissues revealed a shift in the lipidome of MLKL deficient mice on a HFD. Adding to MLKL’s new role in adipocyte differentiation^78^, we discerned a possible role for MLKL in the synthesis of lipotoxic saturated ceramide species in the VAT, which are associated with reduced adipocyte function and global insulin resistance^65^. Secondly, we observed a prominent defect in diglyceride (DG) and triglyceride (TG) production in *Mlkl^-/-^* mice, particularly MUFA and PUFA species, showing a new role for MLKL in lipid biosynthesis. Interestingly, NASH in RIPK3-deficient mice has also been associated with a more select reduction in DG and TG species with longer acyl chains and greater double bonds^26^, which may suggest some functional overlap between RIPK3 and MLKL in the production of long and very long chain fatty acids that may promote necroptosis^63^.

Our study uncovers a role for RIPK3-caspase-8 signaling in regulating obesity-induced inflammation, independent of its function to activate NLRP3, aligning with recent reports in sepsis and arthritis models^31,53^. More importantly, we delineate a noncanonical RIPK3-indepdendent role for MLKL in the development of obesity and NAFLD development. How MLKL is triggered in this scenario remains elusive, but it has been suggested that RIPK1 may target MLKL to drive necroptosis during liver disease^21,28,83^. We also cannot fully exclude a canonical RIPK3-MLKL signaling axis exists, as RIPK3-caspase-8 activity in distinct cell types may obscure necroptotic events *in vivo*. It is also plausible that non-necroptotic MLKL activity may be triggered by an as-yet-unknown event, such as has been posited for demyelinating diseases^84^.

The vital role for MLKL in regulating lipid uptake, transport and metabolism that we show here suggests future studies investigating the proximity of MLKL to specialized metabolic organelles and examination of the expression and function of lipid receptors/transporters, in the absence of MLKL, are warranted. Ultimately, our study also offers a new avenue and perspective on how targeting divergent MLKL functions, beyond cell death, may limit obesity and NAFLD.

## METHODS

### Mice

All mice were housed under standard regulatory conditions at the Walter and Eliza Hall Institute of Medical Research (WEHI), Australia, and Baker Heart and Diabetes Institute, Australia. All procedures were performed in accordance with the National Health and Medical Research Council Australian Code of Practice for the Care and Use of Animals and approved by the WEHI Animal Ethics Committee or the AMREP AEC. Wild-type (WT), MLKL-deficient (*Mlkl^-/-^*)^17^, RIPK3-deficient (*Ripk3^-/-^*)^85^, RIPK3/Caspase-8 doubly-deficient (*Ripk3^-/-^Casp8^-/-^*)^86^, Caspase-1-deficient (*Casp1^-/-^*)^87^ and NLRP3-deficient (*Nlrp3^-/-^*)^88^ mice, generated or backcrossed onto the C57BL/6J background, were used for the *in vitro* generation of bone marrow-derived macrophages (BMDMs) at > 6 weeks of age. For the *in vivo* high-fat diet model, WT mice harbouring a floxed caspase-8 allele (*Casp8^lox/lox^*) were first crossed onto a RIPK3-deficient background (*Casp8^lox/lox^Ripk3^-/-^*). These mice were then used to generate mice with a conditional deletion of caspase-8 in myeloid cells using the Lysozyme M Cre transgenic mouse (*Casp8^LysMCre^Ripk3^-/-^*). To obtain optimal numbers of age-matched male mice of relevant mouse lines the following breeding strategies were adopted and genotypes pooled by post-natal day 35 and acclimatized for at least 3 weeks. *Casp8^lox/lox^* (or *Casp8^LysMCre/+^*) mice were generated using *Casp8^lox/lox^* x *Casp8^lox/lox^* and/or *Casp8^LysMCre/+^* x *Casp8^lox/lox^* crosses. *Casp8^lox/lox^Ripk3^-/-^* and *Casp8^LysMCre^Ripk3^-/-^* mice were obtained in parallel mating from *Casp8^lox/lox^Ripk3^-/-^* x *Casp8^LysMCre^Ripk3^-/-^* crosses. WT and *Mlkl^-/-^* mice were generated from heterozygous and/or homozygous *Mlkl^-/-^* and WT *Mlkl^+/+^* matings. Bone marrow chimeric mice were generated by irradiating C57BL/6 recipient mice twice with doses of 5.5 Gg, spaced 3 h apart, and intravenously injecting 5 x10^6^ Ly5.2 WT or *Mlkl^-/-^* donor bone marrow cells (post red blood cell lysis) *via* the tail vein in 200 μL PBS. Mice were allowed to reconstitute for 8 weeks prior to dietary challenge.

### Diets

Eight-to nine-week old WT, *Mlkl^-/-^, Casp8^lox/lox^, Casp8^LysMCre/+^, Casp8^lox/lox^Ripk3^-/-^* and *Casp8^LysMCre^Ripk3^-/-^* mice were fed either a high-fat diet (HFD; 36% fat, 59% of total energy from lipid; Specialty Feeds, WA, Australia) or a normal chow diet (ND) *ad libitum* for 16-26 weeks, as performed previously^89^. Cohorts were weighed weekly in order to measure weight gain. HFD-fed mice (irrespective of genotype) that did not achieve a 25% weight gain by 25 weeks were excluded from further analysis (*i.e*., deemed non-responders), as were animals that developed malocclusion. In the case of BM chimeras, mouse body composition (lean and fat mass) was measured using a 4-in-1 EchoMRI body composition analyzer (Colombus Instruments). Food input and output were grossly monitored by the amount of food consumed and cage weight on a weekly basis. At specified times, or at the experimental endpoint, blood was collected *via* cardiac bleed for serum collection by centrifugation. Organs and tissues, including the liver, spleen, kidney, pancreas, subcutaneous adipose tissue (SAT) and visceral adipose tissue (VAT) were harvested and weights recorded prior to further analysis.

### Glucose and insulin tolerance tests

Intraperitoneal (IP) glucose tolerance tests were performed in ND and HFD challenged mice at 8-10 weeks and 16-18 weeks, and an insulin tolerance test was performed after 23 weeks of diet challenge. In both cases, mice were fasted for 5-6 h before being given an intraperitoneal injection of either 1.5 g D-glucose per kg of body weight or 0.75 U insulin per kg of body weight. Oral glucose tolerance tests (2 g/kg by oral gavage) were performed on bone marrow chimeric mice based on lean body mass. Blood glucose levels were measured (Accu-Chek Performa, Roche) by tail bleeds prior to injection or oral gavage (time 0 min), and measurements made at 15, 30, 50, 80 (90) and 120 min post-glucose or insulin administration.

### Serum analysis

Serum triglycerides, cholesterol, glucose, alanine aminotransferase (ALT) and aspartate aminotransferase (AST) levels were measured by ASAP Laboratory (Mulgrave, VIC, Australia). Insulin and NEFA levels were using a mouse insulin ELISA kit (Promega) and WAKO NEFA-C kit (WAKO) according to the manufacturer’s instructions.

### Histopathology

Liver and VAT biopsies were fixed in 10% (w/v) neutral-buffered formalin and blocked in paraffin. Sections (4 μm) were stained with Haematoxylin and Eosin (H&E), Periodic-Acid Schiff (PAS), Sirius red (SR), or immunostained for F4/80 antibody (in-house; WEHI histology services). Liver samples were also snap-frozen in Tissue-tek OCT and 8 μM sections cut before Oil Red O staining to illustrate lipid content. Liver inflammation, steatosis and ballooning were scored on H&E (or PAS) stained sections, and fibrosis assessed on Sirius Red stained section, by an independent pathologist that was blinded to the study details (Gribbles Veterinary Pathology, Clayton, VIC, Australia) using the NASH-Clinical Research Network criteria. An automated script was created in FIJI (WEHI Centre for Dynamic Imaging; available from the authors on request) that used a MorphoLibJ plugin to segment adipocytes for quantification of the mean VAT adipocyte size^90,91^. Adipocyte measurements were performed in 2-4 focal regions per H&E stained tissue section. Areas with < 100 adipocytes within quadrant and quadrants with poor tissue integrity for quantification were excluded from the analysis.

### Flow cytometric analysis

VAT and liver tissue (0.5-1 g) was harvested from mice and minced into small pieces (3-4 mm) with surgical scissors, and then enzymatically dissociated in 1 mg/mL Type I collagenase (Worthington) and DNAse 1 (5 ng/mL) in 2% (v/v) FBS in DME for 45 min at 37°C (vortexing every 5 min) before adding 2.5 mM EDTA for final 15 min. Cells were sieved through a 70 μm cell strainer (Falcon), washed in 3% (v/v) FBS containing 2.5 mM EDTA in PBS (FACS buffer) and leukocytes pelleted at x 800 G for 15-20 min with the brake off. Red blood cells were lysed, cells washed and resuspended in FACS buffer and subsequently stained with fluorochrome-conjugated antibodies from Biolegend, BD Bioscience and Ebioscience to mouse CD16/32 (Fc block, 2.4G2), CD45.2 (104), CD11b (Mac-1), F4/80 (BM1), Ly6G (1A8), Ly6C (HK1.4), CD11c (N418; in-house) and MHCII (M5/114.15.2; in-house) for 30 min on ice. Cells were washed and resuspended in FACS buffer containing propidium iodide (PI, 1 μg/mL) and counting beads (123count eBeads™; Invitrogen) prior to analysis on an LSR-Fortessa instrument using FACSDiva software, and analysis using WEASEL software version 2.7/2.8 (purchased from Frank Battye).

### Preparation of bovine-serum albumin (BSA)-conjugated palmitate (PA)

In order to obtain a 5:1 molar ratio of palmitate (PA):bovine serum albumin (BSA), 1% (w/v) fatty acid-free BSA (Worthington, USA) was prepared by dissolving fatty acid-free BSA in serum-free DMEM containing 4 μM L-glutamine. Stocks of 150 mM PA were prepared by dissolving 41.8 mg sodium palmitate (Sigma-Aldrich) in 1 mL 50 % ethanol at 70 °C for 5 mins. BSA was pre-incubated at 37 °C for 30 min before conjugation to PA for 1 h at 37 °C.

### Bone marrow derived macrophage cultures

Bone marrow cells were harvested from the tibial and femoral bones to generate BMDMs. Cells were cultured in DMEM media (Gibco) containing 10% (v/v) foetal bovine serum (FBS; Bovogen), 15-20% (v/v) L929-conditioned media, 4 μM L-glutamine (Life Technologies), 1 mM sodium pyruvate (Thermofisher), and 100 U/mL penicillin/streptomycin (P/S) (Life Technologies) for 6 days at 37 °C, 10% CO_2_. Unless otherwise indicated, macrophages were plated at 4 × 10^5^ cells/well in 24-well tissue culture-treated plates (BD Falcon), or at 3 × 10^5^ in 24-well non-tissue culture-treated plates. Macrophages were primed with B4 or B5 LPS (both 50 ng/mL Ultrapure, Invivogen) or Pam3CSK4 (500 ng/mL, Invivogen) for 3 h before addition of 300-1200 μM of fatty acid free bovine serum albumin conjugated to palmitate (BSA-PA) or BSA-only (to match the maximum BSA added in palmitate stimulations). After 18 h, cell supernatants were routinely collected from tissue culture for cytokine analysis by ELISA and supernatants and cell lysates collected for immunoblotting. In some cases, cells were harvested from non-tissue culture treated plates using 5 mM EDTA in phosphate buffered saline (PBS) to lift adherent cells. Cell viability was measured *via* PI (1-2 μg/mL) uptake and flow cytometric analysis on an BD LSR-Fortessa X20 or BD FACSCanto instrument using FACSDiva software (BD Biosciences). Data were analyzed using FlowJo software version 10.6.1 or WEASEL software.

### Ex vivo VAT cultures

VAT tissue (0.5 g) was harvested from ND and HFD challenged mice and cultured in a well of a 24-well tissue culture plate in 0.1% (w/v) BSA/DMEM in the presence or absence of LPS (50 ng/ml) for 16-18 h at 37 °C, 10% CO_2_. Supernatants were collected for cytokine analysis.

### Cytokine analysis

IL-1β (R&D) and TNF (eBioscience) ELISA kits were used, according to the manufacturers’ instructions. For detection of TNF in cell supernatants, samples were diluted 1:10 in assay diluent.

### Immunoblotting

Cell lysates and supernatants were boiled for 10 min in 1× NuPAGE LDS (Thermofisher) or in-house (2 % w/v SDS, 10 % v/v glycerol, 50 mM Tris pH 6.8, 0.01 % bromophenol blue) sample buffer containing 5 % β-mercaptoethanol (β-ME). Samples were separated on 4-12% Bis-Tris gradient gels (Invitrogen, NW04125/27BOX) and proteins were transferred onto nitrocellulose membrane (Millipore). Ponceau staining was used to confirm protein transfer and as a loading control. Membranes were blocked with 5% (w/v) skim milk in Tris-Buffered Saline (TBS) containing 0.1% (v/v) Tween-20 (TBS-T) for 1 h and then probed overnight at 4°C with the following primary antibodies (all diluted 1:1000 5% (w/v) skim milk in TBS-T containing + 0.02 % sodium azide, with the exception of β-actin that was diluted 1:5000): pro- and cleaved IL-1β (R&D; AF-401-NA), pro- and cleaved caspase-1 (Adipogen; AG-20B-0042-C100), pro-caspase-8 (WEHI; 3B10), cleaved caspase-8 (CST; 9429S), NLRP3 (Adipogen, AG-20B-00140-C100), β-actin (CST; 5125S and 13E5). Relevant horseradish peroxidase (HRP)-conjugated primary and secondary antibodies (all diluted 1:5,000) were applied for 1 h at RT in 5% (w/v) skim milk in TBS-T. Membranes were washed 6x in TBS-T between each incubation. Membranes were developed using the Immobilon Forte Western HRP substrate (Merck Millipore, WBLUF0500) and imaged with a BioRad ChemiDoc MP. Images were analyzed and processed with BioRad ImageLab software.

Snap-frozen liver (100 mg) tissue was ground using a mortar and pestle on dry ice and tissue homogenized (by pipetting) in 600 μL RIPA buffer [150 mM NaCl, 50 mM Tris (pH 7.4), 1 mM deoxycholate, 1% (v/v) Triton-X containing cOmplete™ protease inhibitor cocktail (Roche) and PhosSTOP (Roche)] and agitated on a rotating wheel for 1 hr at 4°C. Following centrifugation (x 20,000 G) at 4°C for 15 min the lipid layer was discarded, and the supernatant transferred to a fresh tube. This process was repeated two more times. Tissue lysate protein concentration was quantified using the DC protein assay (BioRad) according to the manufacturer’s instructions, and 40 μg of tissue lysate was analyzed by immunoblotting, as above, with goat anti-CD36 Affinity Purified polyclonal antibody (SR-B3, R&D Systems, In vitro Life Science) and β-actin, and relevant secondaries.

### Quantitative Polymerase chain reaction (qPCR)

Liver and VAT biopsies were snap-frozen and 50 mg and 100 mg of tissue, respectively, was ground in a mortar and pestle over dry ice. RNA was extracted using Trizol RNA Isolation Reagents (Life Technologies) and on-column DNase treatment and purification were performed using the ISOLATE II RNA mini kit (Bioline) or RNeasy Mini kit (Qiagen) according to the manufacturer’s instructions. RNA concentration and purity were quantified using the Nanodrop 2000 Spectrophotometer, and cDNA synthesis was performed using a High Capacity cDNA synthesis kit (Applied Biosystems, ThermoFisher Scientific). qPCR was then performed using the Power SYBR™ green PCR Master Mix (Applied Biosystems) on a QuantStudio 6 Flex PCR system (ThermoFisher Scientific) with the primer pairs listed in Supplementary Table 1. Relative mRNA levels were calculated using the comparative delta CT (dCT) method (2^-[(dCT genotype/diet) – (dCT WT ND)]^), where dCT values were obtained by normalization to the internal house-keeping reference gene *18s*. Specificity of each primer set was confirmed by the observation of a single peak in the melt curve graph of each qPCR run.

### 3′ mRNA sequencing

RNA from liver samples were extracted as above. Integrity of RNA was examined using Tapestation Agilent 4200 and samples with RINe>8 were selected for library preparation for 3’ mRNA-sequencing analysis. 3’mRNA-sequencing libraries were prepared using 100 ng of total RNA using the QuantSeq 3’mRNA-Seq Library Prep (Lexogen) according to the manufacturer’s instructions. The single-end 75 bp were demultiplexed using CASAVAv1.8.2 and Cutadapt (v1.9) was used for read trimming^92^. The trimmed reads were subsequently mapped to the mouse genome (mm10) using HISAT2^93^. FeatureCounts from the Rsubread package (version 1.34.7) was used for read counting after which genes without a counts per million reads (CPM) in at least 3 samples were excluded from downstream analysis^94,95^. Count data were normalized using the trimmed mean of M-values (TMM) method and differential gene expression analysis was performed using the limma-voom pipeline (limma version 3.40.6)^94,96,97^. Comparisons between WT ND, *Mlkl^-/-^* ND, WT HFD and *Mlkl^-/-^* HFD were made. GSEA 2.2.2 was used for Gene set enrichment analysis (GSEA)^98,99^. Gene ontology (GO) and analysis was performed using *Metascape*, and *pheatmap* (version 1.0.12) was used to generate heatmaps. The datasets generated during this study are available at GEO225560.

### Lipidomic analysis

Targeted lipidomic analysis using Liquid Chromatography-Mass Spectrometry (LC-MS) was performed on the serum, liver and VAT of ND- and HFD-fed WT and *Mlkl^-/-^* mice. Briefly, for serum, 20 μL was aliquoted (serum from up to 3 mice was pooled where necessary) into Eppendorf tubes and 380 μL of cold 2:1 chloroform/methanol containing 4 internal standards (10 mg/L; PG 17:0/17:0, PC 19:0/19:0, PE-D31 16:0/18:1 and TG-D5 19:0/12:0/19:0 (Product # 860374, 850367, 8609040 and 830456 from Avanti Polar lipids, Alabama, USA) were added. For liver and VAT lipid extraction, snap-frozen tissue (~ 30 mg) was transferred into cryomill tubes and 500 μl of cold 1:9 chloroform: methanol (v/v) containing 4 internal standards as mentioned earlier (10 mg/L) were added before homogenisation using a cryogenically cooled bead-mill (6,800 rpm, 3 x 45 sec cycles; Precellys24 coupled to Cryolys unit from Bertin technologies). Next, 400 μL of homogenate was transferred into Eppendorf tubes and 680 μL of 100% methanol was added to make final concentration of 2:1 chloroform: methanol (v/v). Serum, VAT and liver samples were then vortexed (30 sec), mixed with a thermomixer (Eppendorf South Pacific Pty Ltd, Macquarie Park, Australia) at 950 rpm for 10 min at 20°C and centrifuged at 15,000 rpm (Beckman Coulter Microfuge^®^ 22R refrigerated microcentrifuge, Beckman Coulter Australia Pty Ltd, Sydney, Australia) for 10 min at room temperature. 1250 μL of VAT and liver or 350 μL of serum supernatant was then transferred into Eppendorf tubes containing glass inserts and gradually evaporated (50 μL at a time) using a vacuum concentrator, with a temperature maintained at 30-35°C (Christ^®^ RVC 2-33, Martin Christ Gefriertrocknungsanlagen GmbH, Osterode am Harz, Germany). Samples were reconstituted with 10μL methanol: 90μL water saturated butanol (100μL, v/v) for LC-MS analysis. Pooled biological quality controls (PBQC) were prepared by pooling aliquots of the extracts from each sample and were run after every five samples.

Extracted lipids were processed and detected by Metabolomics Australia (Bio21 Institute, Melbourne, Australia) using an Agilent 1290 LC system and Agilent Triple Quadrupole 6490 mass spectrometer (Agilent Technologies Australia, Mulgrave, Australia), as previously described^100^. Briefly, lipids from the serum, VAT and liver samples were separated using a Zorbax Eclipse Plus C18 column (100mm x 2.1 mm x 1.8um, Agilent technologies Australia). Injection volume was kept at 1uL with a LC flowrate of 400 uL/min. The LC mobile phase solvents were acetonitrile/water/isopropanol (30:50:20, v/v/v) for mobile phase A and acetonitrile/water/isopropanol (9:1:90, v/v/v) for mobile phase B with 10mM ammonium formate for both A and B. The LC gradient, MS parameters and the targeted dynamic scheduled multiple reaction monitoring (dMRM) transitions of each lipid species have been previously described^100,101^. Quantitation was based on relative changes in peak areas. Data processing was performed using Agilent’s Mass Hunter Quantitative Analysis software (Agilent Technologies Australia). Lipids were named according to nomenclature described in LIPID MAPS, by Liebisch et al^102^.

For analysis, raw data was normalized for median lipid content per mouse and the weight of tissue analyzed, as appropriate. These data were subjected to a log10 transformation for statistical analyses. Total Relative abundance of lipid specie classes were calculated by summation of individual normalized species, prior to log10 transformation for statistical analyses (mean ± SEM). The Log2 Fold change was also calculated in Metaboanalyst 5.0 from the median and weight normalized HFD sample data and adjusted for a false discovery rate (FDR) to detect significantly different lipid species in the serum and liver (p < 0.05, T-test). Heatmaps where generated using GraphPad PRISM (Version 9) software and are presented as median abundance of lipid species. The raw data and internal quality control samples are available on request.

### Statistical analysis

All graphical data are presented as the mean ± standard error of the mean (SEM) for biological samples or standard deviation (SD) for replicates, as indicated in the figure legends. AUC was calculated for weights, % weight gain and GTT and ITT tests. Statistical comparisons between two genotypes was performed using a Student’s t-test and three genotypes a one-way analysis of variance (ANOVA) was performed. A two-way ANOVA with a Tukey *post-hoc* correction for multiple comparisons was used for comparisons between treatments and multiple genotypes. Analyses were performed using GraphPad PRISM (Version 9) software and a *p* value of *p* < 0.05 was considered statistically significant.

## Author Contributions

The project was conceived by J.E.V. and K.E.L. The experiments were designed by S.A.C., H.T., H.K., I.K., V.K.N., D.P.D., E.D.H., A.J.M., J.E.V. and K.E.L., and performed by S.A.C., H.T., T.M.D., N.A., H.K., I.K., M.S., J.E., D.S.S., C.H., A.J.V., V.K.N., S.R., R.C., M.R., L.W., J.E.V. and K.E.L. The manuscript written by S.A.C., H.T., T.M.D., M.S., T.A.G., J.E.V. and K.E.L. Expert advice and essential mice and reagents were provided by J.M.H., J.M.M., D.P.D., S.L.M., E.D.H. and A.J.M. All authors assisted with data interpretation and manuscript editing.

## Acknowledgements

We thank WEHI Bioservices and Alfred Research Alliance Animal Facility for expert animal care, WEHI’s histology laboratory and the Centre for Dynamic Imaging for histopathology staining and microscopy assistance, respectively. We thank WEHI and Monash FlowCore for assistance with FACS. We thank Professor John Silke, Professor Terry Speed (WEHI) and Dr Najoua Lalaoui (Peter MacCallum Cancer Centre) for reagents and helpful advice. We gratefully acknowledge grant support from the National Health and Medical Research Council (NHMRC) of Australia: project grants (1140187, 1165591 to E.D.H.; 1145788 to J.E.V., K.E.L. and J.M.M.; 1101405 to J.E.V.; 1162765 to K.E.L.), Ideas grants (2011584 to M.S. and J.M.H.; 1183070 to J.E.V.; 1181089 to K.E.L.) and fellowships (CJ Martin Overseas Biomedical Training Fellowship 1144014 to S.A.C.; Leadership Fellowship; 1172929 to J.M.M.; 2008652 to E.D.H.; 1194329 to A.J.M.; 1141466 to J.E.V.). K.E.L. is an Australian Research Council (ARC) Future Fellow (FT190100266). This work was also supported by operational infrastructure grants through the Australian Government Independent Research Institute Infrastructure Support Scheme (9000719) and the Victorian State Government Operational Infrastructure Support, Australia.

