## Supplemental Figures and Table for "Divergent roles for caspase-8 and MLKL in high-fat diet induced obesity and NAFLD in mice"

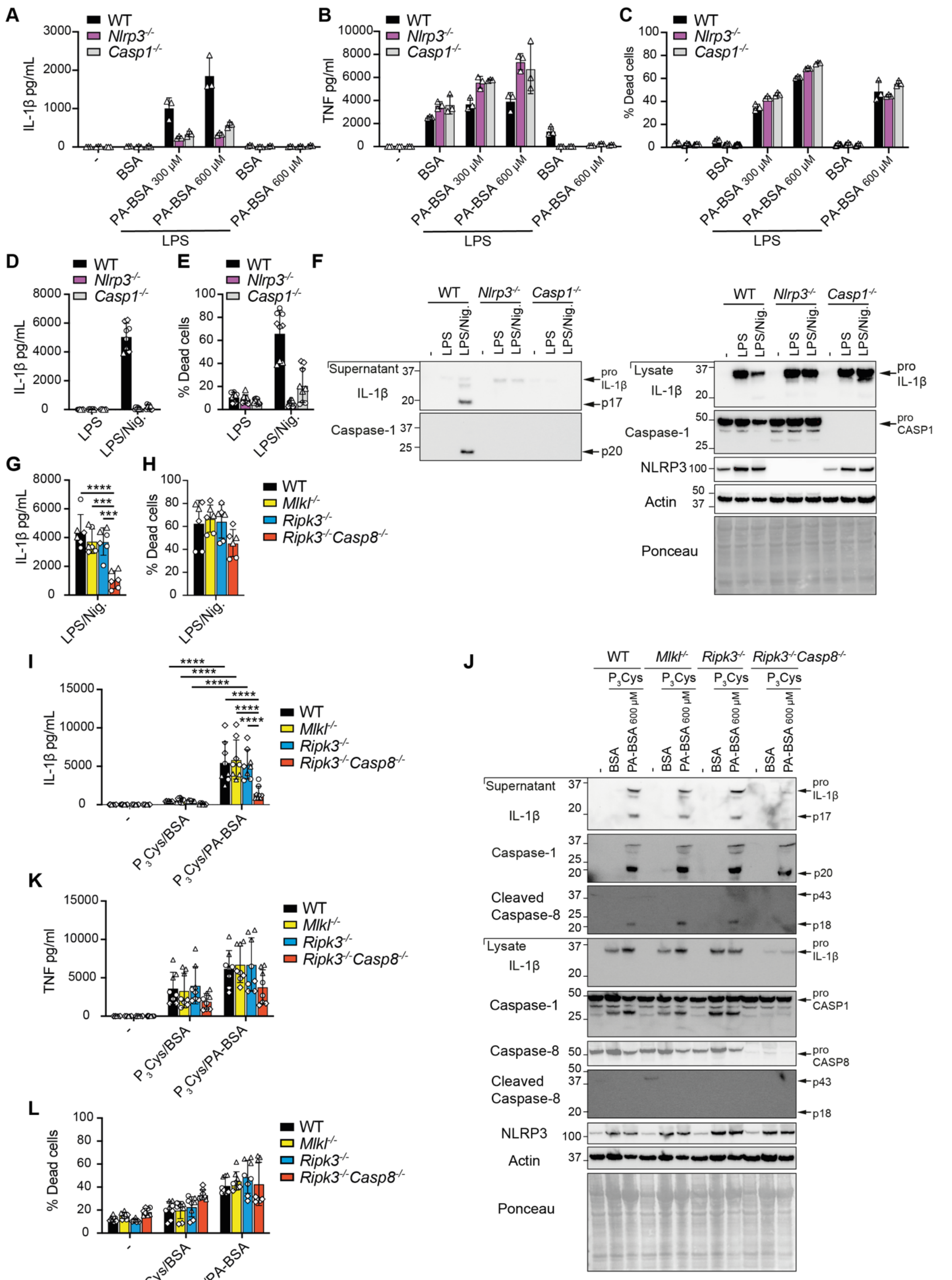

A
WT
Nlrp3-/-
Casp1-/-
IL-1β pg/mL
3000
2000
1000
0
-
BSA
PA-BSA 300 μM
PA-BSA 600 μM
BSA
PA-BSA 600 μM
LPS
B
TNF pg/ml
10000
8000
6000
4000
2000
0
C
% Dead cells
100
80
60
40
20
0
D
8000
6000
4000
2000
LPS
LPS/Nig.
E
F
Supernatant
IL-1β
Caspase-1
pro IL-1β
p17
p20
Lysate
pro CASP1
NLRP3
Actin
Ponceau
G
****
***
H
Mlkl-/-
Ripk3-/-
Ripk3-/-Casp8-/-
I
15000
10000
5000
P3Cys/BSA
P3Cys/PA-BSA
J
P3Cys
Cleaved Caspase-8
p43
p18
pro CASP8
K
L


Figure S1

**Supplementary Fig. 1 Defective caspase-8-mediated inflammasome priming reduces palmitate-induced NLRP3 inflammasome activation.**

**(A-C)** WT, *Nlrp3*$^{-/-}$ and *Casp1*$^{-/-}$ BMDMs were pre-treated with and without LPS (50 ng/ml) for 3 hours and treated with 300-600 μM palmitate conjugated to BSA (PA-BSA) or BSA alone (equivalent to 600 μM BSA amount) for a further 18-20 h. **(A)** IL-1β and **(B)** TNF levels were measured in cell supernatants by ELISA and **(C)** cell viability was assessed by PI incorporation and flow cytometric analysis. (n = 3 replicates). **(D-F)** WT, *Nlrp3*$^{-/-}$ and *Casp1*$^{-/-}$ BMDMs were primed with LPS (50 ng/ml) for 3 h and treated with nigericin (10 μM) for ~ 45 min. **(D)** IL-1β levels were measured in cell supernatants by ELISA, and **(E)** cell viability was assessed by PI incorporation and flow cytometric analysis. (n = 3 replicates pooled from 3 independent experiments). **(F)** Cell lysates and supernatants were analyzed by immunoblot. Immunoblots are representative of one of 2 experiments. **(G, H)** WT, *Mlkl*$^{-/-}$, *Ripk3*$^{-/-}$ and *Ripk3*$^{-/-}$*Casp8*$^{-/-}$ BMDMs were primed with LPS (50 ng/ml) for 3 h and treated with nigericin (10 μM) for ~ 45 min. **(G)** IL-1β levels were measured in cell supernatants by ELISA, and **(H)** cell viability was assessed by PI incorporation and flow cytometric analysis. (n = 2 replicates pooled from 3 independent experiments). **(I-L)** WT, *Mlkl*$^{-/-}$, *Ripk3*$^{-/-}$ and *Ripk3*$^{-/-}$*Casp8*$^{-/-}$ BMDMs were primed with Pam3Cys (500 ng/ml) for 3 h and treated with 600 μM PA-BSA or BSA alone, as indicated, for 18-20 h. **(I)** IL-1β and **(K)** TNF levels were measured in cell supernatants by ELISA. (n = 2-3 replicates pooled from 3 independent experiments). **(J)** Cell lysates and supernatants were analyzed by immunoblot. Data are representative of one of 2 experiments. **(L)** Cell viability was assessed by PI incorporation and flow cytometric analysis. (n = 2-3 replicates pooled from 3 independent experiments). For individual experiments, replicates are shown as different shapes.

Data are the mean ± SD. *p < 0.05; **p < 0.01; ***p < 0.001; ****p < 0.0001. Two-way ANOVA followed by Tukey's multiple comparisons test was applied. See related **Fig. 1.**

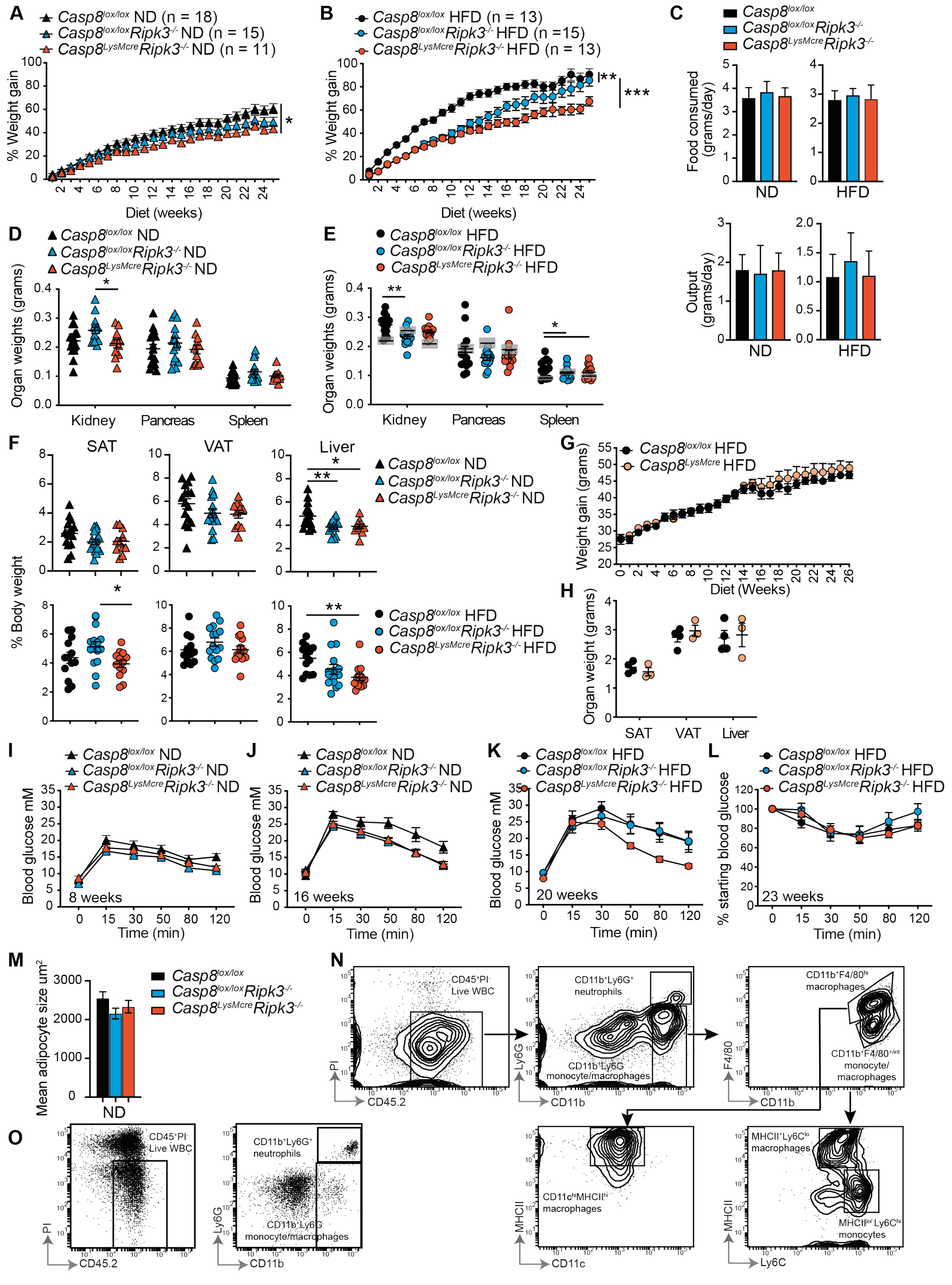
A
Casp8lox/lox ND (n = 18)
Casp8lox/loxRipk3-/- ND (n = 15)
Casp8LysMcreRipk3-/- ND (n = 11)
% Weight gain
Diet (weeks)
B
Casp8lox/lox HFD (n = 13)
Casp8lox/loxRipk3-/- HFD (n =15)
Casp8LysMcreRipk3-/- HFD (n = 13)
C
Casp8lox/lox
Casp8lox/loxRipk3-/-
Casp8LysMcreRipk3-/-
Food consumed (grams/day)
Output (grams/day)
ND
HFD
D
Casp8lox/lox ND
Casp8lox/loxRipk3-/- ND
Casp8LysMcreRipk3-/- ND
Organ weights (grams)
Kidney
Pancreas
Spleen
E
Casp8lox/lox HFD
Casp8lox/loxRipk3-/- HFD
Casp8LysMcreRipk3-/- HFD
F
SAT
VAT
Liver
% Body weight
G
Casp8lox/lox HFD
Casp8LysMcre HFD
Weight gain (grams)
Diet (Weeks)
H
Organ weight (grams)
SAT
VAT
Liver
I
J
K
L
Blood glucose mM
% starting blood glucose
Time (min)
8 weeks
16 weeks
20 weeks
23 weeks
M
Mean adipocyte size um2
N
CD45+PI- Live WBC
CD11b+Ly6G+ neutrophils
CD11b+Ly6G- monocyte/macrophages
CD11b+F4/80hi macrophages
CD11b+F4/80+/int monocyte/ macrophages
CD11chiMHCIIhi macrophages
MHCII+Ly6Clo macrophages
MHCIIlo/-Ly6Chi monocytes
PI
CD45.2
Ly6G
CD11b
F4/80
MHCII
CD11c
Ly6C
O

**Figure S2**

**Supplementary Fig. 2 Mice lacking caspase-8 and RIPK3 in myeloid cells are modestly protected from HFD challenge.**

**(A, B)** Wildtype *Casp8*$^{lox/lox}$, *Casp8*$^{lox/lox}$*Ripk3*$^{-/-}$, and *Casp8*$^{LysMcre}$*Ripk3*$^{-/-}$ were fed a **(A)** normal chow diet (ND) or **(B)** high fat diet (HFD) for 25 weeks from 8-9 weeks of age and % weight gain assessed weekly. (n ≥ 11 mice per group pooled from 3 independent experiments). **(C)** Food intake and output per mouse per day was calculated from weekly measurements. (n ≥ 8 mice per group from 2 pooled independent experiments). **(D-E)** End-stage (25 weeks of diet) organ weights and **(F)** organ weights expressed as a % of body weight. (n ≥ 11 mice/group pooled from 3 independent experiments). **(G, H)** *Casp8*$^{lox/lox}$ and *Casp8*$^{LysMcre}$ mice were fed a HFD for 26 weeks and **(G)** weight gain was assessed weekly and **(H)** organ weights measured at the end of the experiment. (n = 3-4 mice from 1 experiment). **(I-L)** Blood glucose was measured at specified times (8-14 weeks and 16-20 weeks) in ND- and HFD-fed mice during intraperitoneal GTT (1.5 g/kg) and insulin tolerance test (ITT, 0.75 Units/kg), as indicated. (n = 5-6 mice per group). Data are representative of one of 2-3 experiments. **(M)** Mean adipocyte size in VAT of ND-fed wildtype *Casp8*$^{lox/lox}$, *Casp8*$^{lox/lox}$*Ripk3*$^{-/-}$, and *Casp8*$^{LysMcre}$*Ripk3*$^{-/-}$ mice after 25 weeks of diet. n = 6-12 mice pooled from 3 independent experiments. **(N, O)** Gating strategy for analyzing live CD45$^{+}$ PI$^{-}$ myeloid populations in the **(N)** VAT and **(O)** liver of mice fed a HFD for 25 weeks.

Data are the mean ± SEM or mean ± SD for (C). *p < 0.05; **p < 0.01; ***p < 0.001; ****p < 0.0001. One-way ANOVA of AUC (A, B, G, I-L) and One-way ANOVA (C-F, H, M). See related **Fig. 2 and 3.**

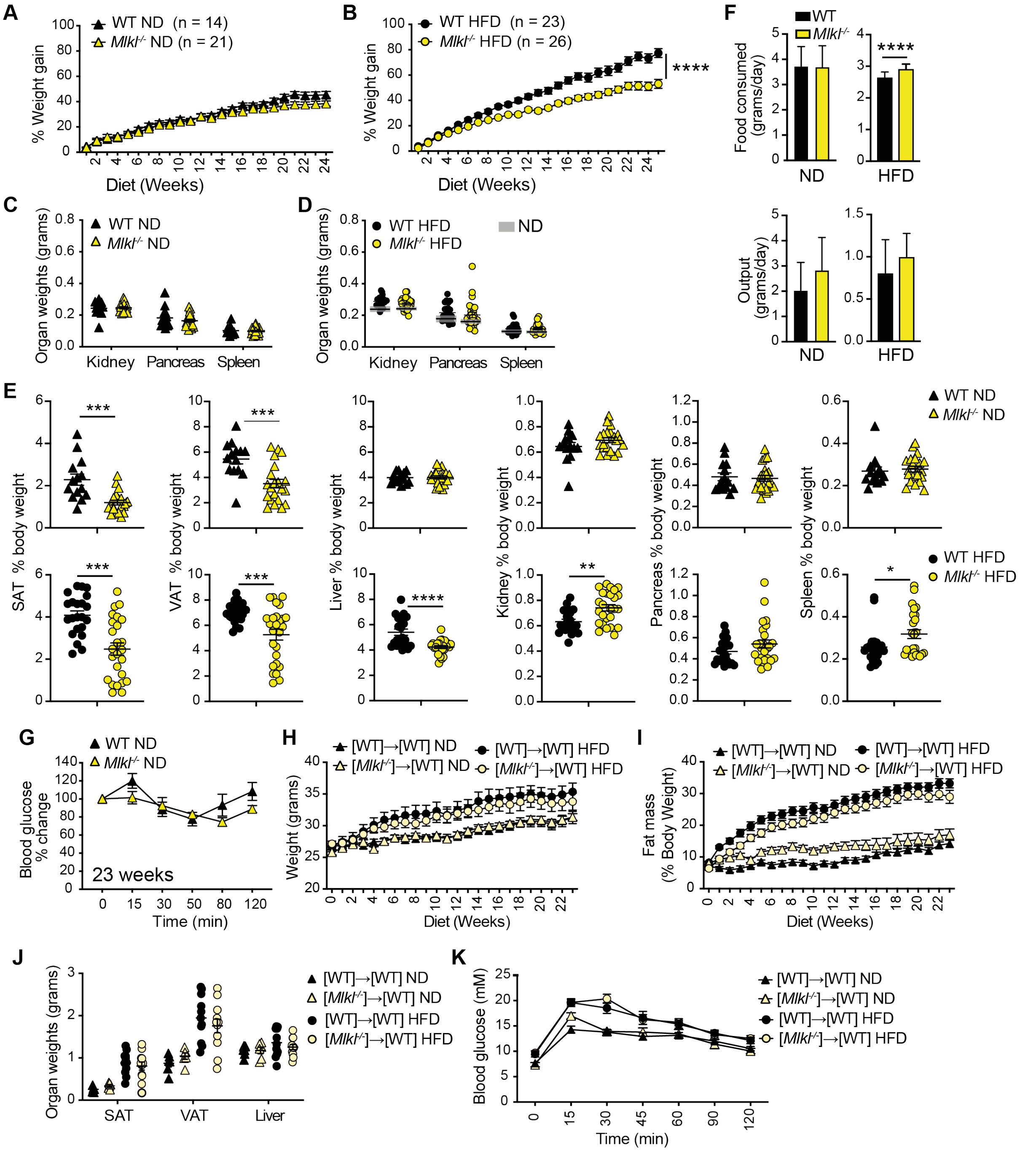


Figure S3

**Supplementary Fig. 3 MLKL deficiency reduces weight gain and adipose tissue expansion.** Wildtype (WT) and *Mlkl*$^{-/-}$ were fed a normal chow diet (ND) or high fat diet (HFD) for 25 weeks from 8-9 weeks of age. **(A, B)** % weight gain from weekly measurements. (ND n ≥ 14 mice/group and HFD n ≥ 23 mice per group pooled from 3 independent experiments). **(C-E)** End-stage organ weights, and **(E)** organ weights expressed as a % of body weight at end of experiment. (n = 11-17 mice per group pooled from 3 independent cohorts). **(F)** Food intake and output. (ND n ≥ 9 mice per group and HFD n ≥ 20 mice/group pooled from 2 independent experiments). **(G)** Blood glucose was measured at specified times in ND mice during insulin tolerance test (ITT, 0.75 Units/kg), as indicated. (n = 5-6 mice per group). **(H-K)** C57BL/6 mice were reconstituted with WT or *Mlkl*$^{-/-}$ bone marrow and fed a ND or HFD for 23 weeks. **(H)** Weight gain and **(U)** fat mass as a % of total body weight assessed using an EcoMRI were measured weekly. **(J)** End-stage organ weights were measured. **(K)** Blood glucose was measured at 10 weeks in ND- and HFD-fed mice following oral gavage of glucose (2 g/kg based on lean body mass). (n = 12 mice per group).

Data are the mean ± SEM or mean ± SD for (F). *p < 0.05; **p < 0.01; ***p < 0.001; ****p < 0.0001. Student's T-test of AUC (A, B, G-I, K) and Student's T-test (C-F, J). See related **Fig. 4 and 5.**

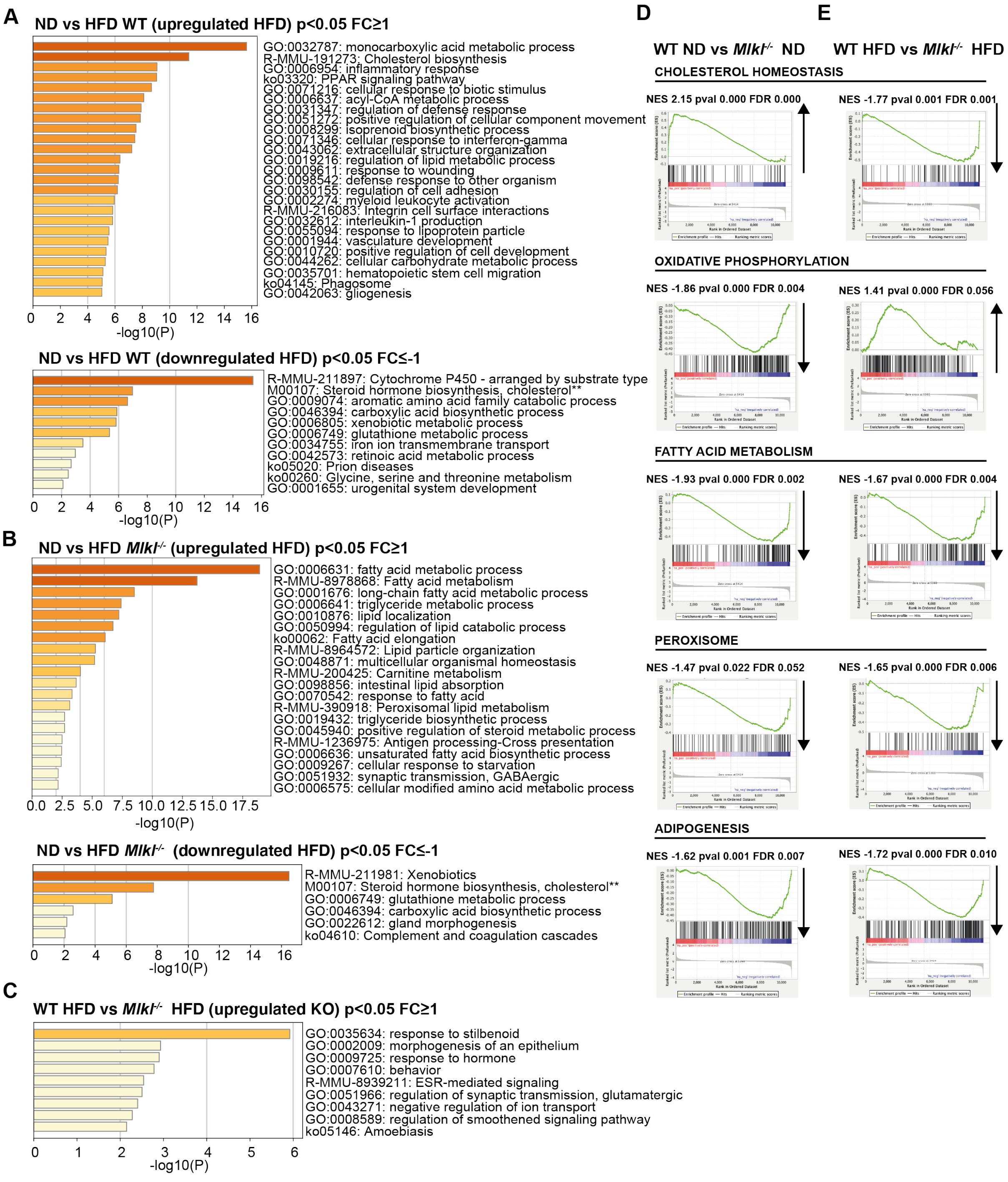


**Figure S4**

**Supplementary Fig. 4 Gene signatures in ageing and HFD challenged MLKL deficient mouse livers.**

Wildtype (WT) and *Mlkl*$^{-/-}$ were fed a normal chow diet (ND) or high fat diet (HFD) for 25 weeks. 3' mRNAseq analysis was performed on the liver samples of 3 mice per diet. **(A, B, C)** Gene Ontology (GO) pathways of significant DEGs upregulated and downregulated (as indicated) in

**(A)** WT HFD-fed versus WT ND-fed, **(B)** ND-fed *Mlkl*$^{-/-}$ livers versus HFD-fed *Mlkl*$^{-/-}$ and **(C)** WT HFD-fed versus *Mlkl*$^{-/-}$ HFD-fed mouse livers. Cut-off values. $p \leq 0.05$ and $logFC \geq 1$ or $logFC \leq -1$. **(D, E)** Gene Set Enrichment analysis (GSEA) enrichment plots for genes differentially regulated between WT versus *Mlkl*$^{-/-}$ on a ND or HFD. A positive normalized enrichment score (NES) (upward arrow) indicates enrichment, whereas a negative NES (downward arrow) indicates downregulation of a specific pathway in *Mlkl*$^{-/-}$ versus WT mice.

Related to **Fig. 6** and **Supplementary Fig. 5**.

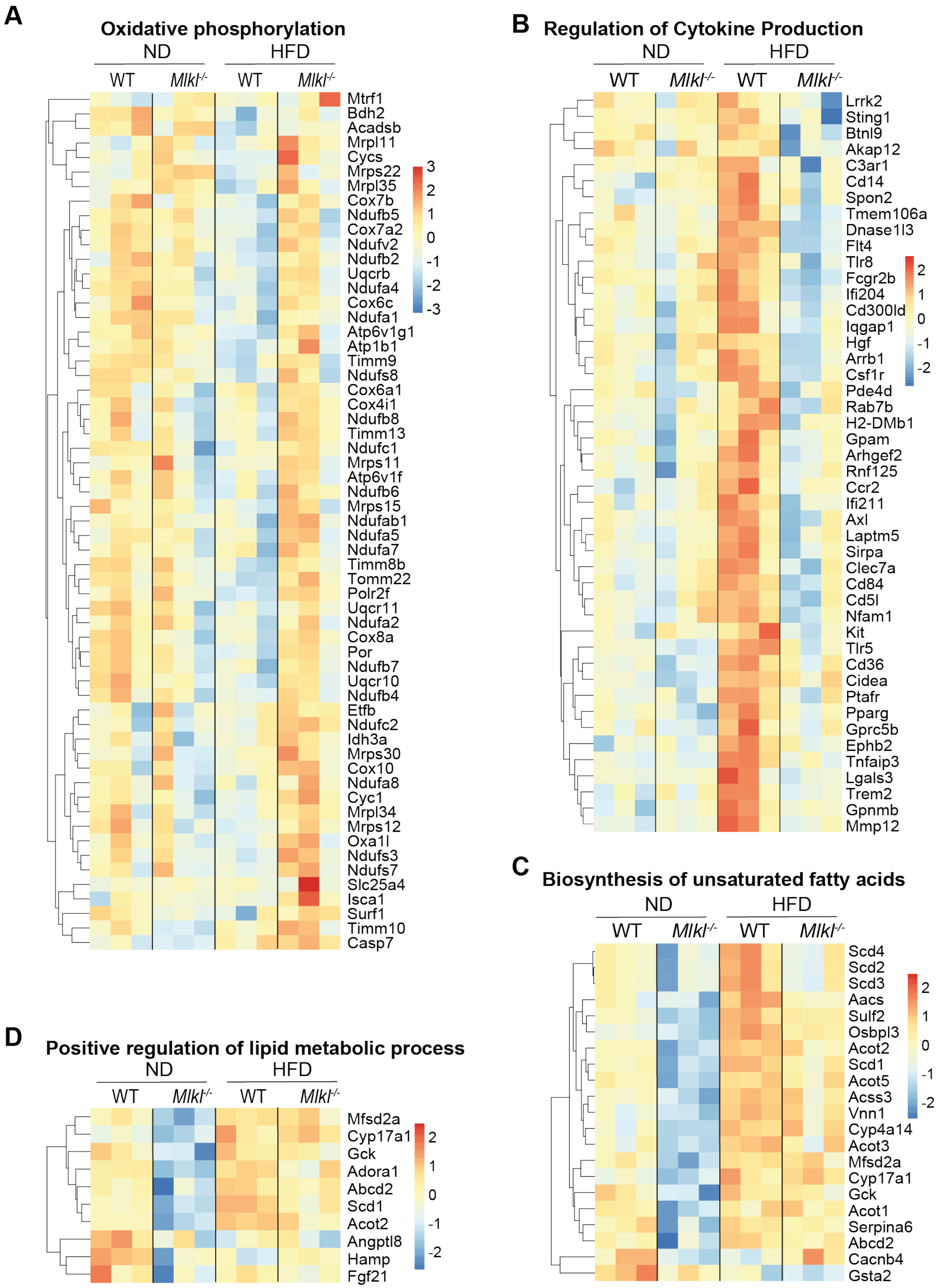
A
Oxidative phosphorylation
ND
HFD
WT
Mlkl-/-
WT
Mlkl-/-
Mtrf1
Bdh2
Acadsb
Mrpl11
Cycs
Mrps22
Mrpl35
Cox7b
Ndufb5
Cox7a2
Ndufv2
Ndufb2
Uqcrb
Ndufa4
Cox6c
Ndufa1
Atp6v1g1
Atp1b1
Timm9
Ndufs8
Cox6a1
Cox4i1
Ndufb8
Timm13
Ndufc1
Mrps11
Atp6v1f
Ndufb6
Mrps15
Ndufab1
Ndufa5
Ndufa7
Timm8b
Tomm22
Polr2f
Uqcr11
Ndufa2
Cox8a
Por
Ndufb7
Uqcr10
Ndufb4
Etfb
Ndufc2
Idh3a
Mrps30
Cox10
Ndufa8
Cyc1
Mrpl34
Mrps12
Oxa1l
Ndufs3
Ndufs7
Slc25a4
Isca1
Surf1
Timm10
Casp7
3
2
1
0
-1
-2
-3
B
Regulation of Cytokine Production
ND
HFD
WT
Mlkl-/-
WT
Mlkl-/-
Lrrk2
Sting1
Btnl9
Akap12
C3ar1
Cd14
Spon2
Tmem106a
Dnase1l3
Flt4
Tlr8
Fcgr2b
Ifi204
Cd300ld
Iqgap1
Hgf
Arrb1
Csf1r
Pde4d
Rab7b
H2-DMb1
Gpam
Arhgef2
Rnf125
Ccr2
Ifi211
Axl
Laptm5
Sirpa
Clec7a
Cd84
Cd5l
Nfam1
Kit
Tlr5
Cd36
Cidea
Ptafr
Pparg
Gprc5b
Ephb2
Tnfaip3
Lgals3
Trem2
Gpnmb
Mmp12
2
1
0
-1
-2
C
Biosynthesis of unsaturated fatty acids
ND
HFD
WT
Mlkl-/-
WT
Mlkl-/-
Scd4
Scd2
Scd3
Aacs
Sulf2
Osbpl3
Acot2
Scd1
Acot5
Acss3
Vnn1
Cyp4a14
Acot3
Mfsd2a
Cyp17a1
Gck
Acot1
Serpina6
Abcd2
Cacnb4
Gsta2
2
1
0
-1
-2
D
Positive regulation of lipid metabolic process
ND
HFD
WT
Mlkl-/-
WT
Mlkl-/-
Mfsd2a
Cyp17a1
Gck
Adora1
Abcd2
Scd1
Acot2
Angptl8
Hamp
Fgf21
2
1
0
-1
-2

**Figure S5**

**Supplementary Fig. 5 Altered gene signatures in MLKL deficient mice in ageing mice and upon HFD.**

Wildtype (WT) and *Mlkl*$^{-/-}$ mice were fed a normal chow diet (ND) or high fat diet (HFD) for 25 weeks. mRNA was extracted from liver tissue (n = 3 mice/group) and 3' mRNAseq analysis performed. **(A)** Heatmap of oxidative phosphorylation genes significantly upregulated in *Mlkl*$^{-/-}$ HFD livers versus WT HFD livers. Cut-off values adjusted $p \leq 0.05$ and $logFC \leq -1$. **(B-D)** Heatmaps of significant DEGs of GO terms **(B)** regulation of cytokine production, **(C)** biosynthesis of unsaturated fatty acids, and **(D)** positive regulation of lipid metabolic process. Cut-off values $p \leq 0.05$ and $logFC \leq -1$.

Related to **Fig. 6 and Supplementary Fig. 4**.

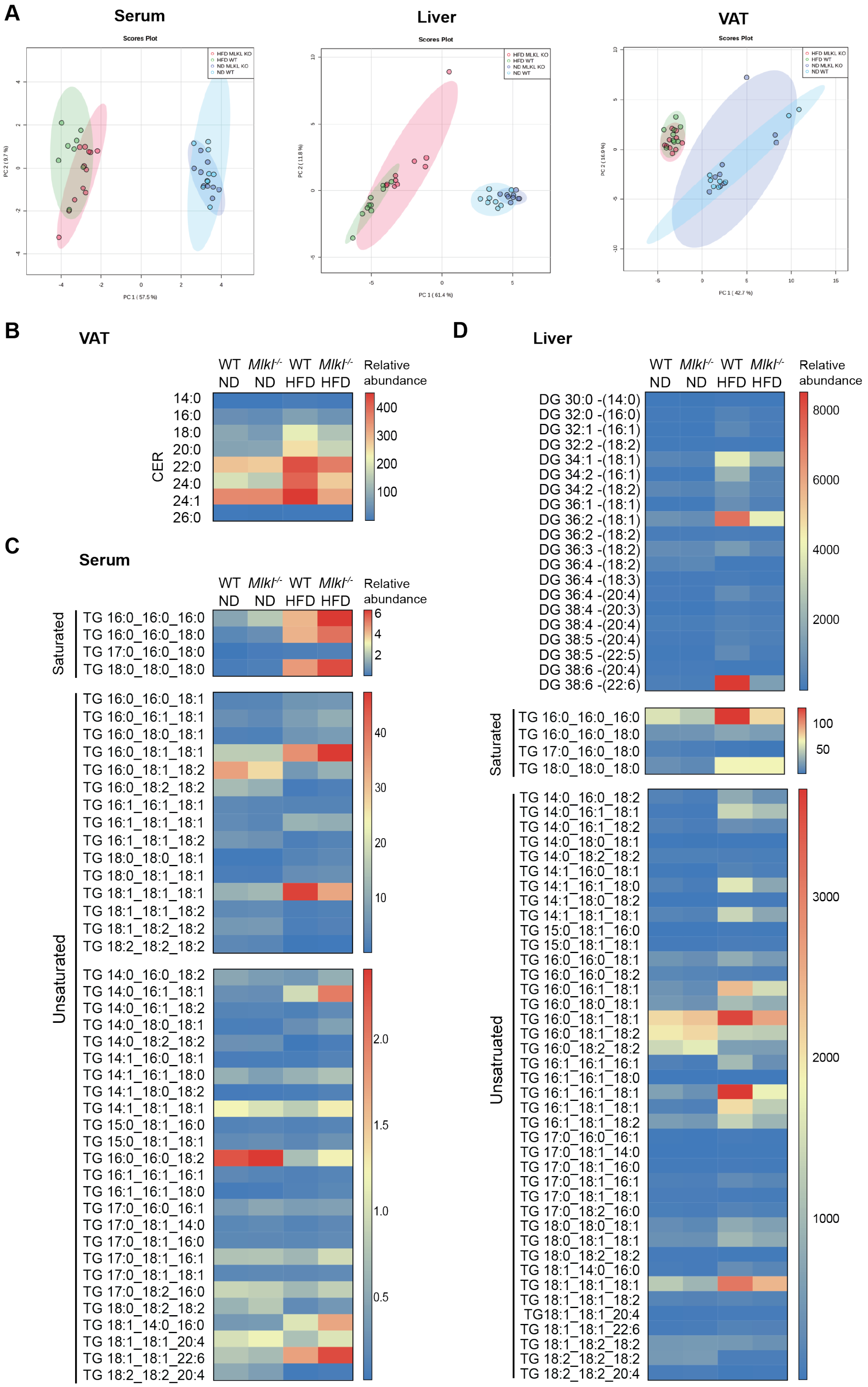

A
Serum
Liver
VAT
Scores Plot
HFD MLKL KO
HFD WT
ND MLKL KO
ND WT
B
VAT
WT ND
Mlkl-/- ND
WT HFD
Mlkl-/- HFD
Relative abundance
CER
14:0
16:0
18:0
20:0
22:0
24:0
24:1
26:0
400
300
200
100
C
Serum
Saturated
TG 16:0_16:0_16:0
TG 16:0_16:0_18:0
TG 17:0_16:0_18:0
TG 18:0_18:0_18:0
6
4
2
Unsaturated
TG 16:0_16:0_18:1
TG 16:0_16:1_18:1
TG 16:0_18:0_18:1
TG 16:0_18:1_18:1
TG 16:0_18:1_18:2
TG 16:0_18:2_18:2
TG 16:1_16:1_18:1
TG 16:1_18:1_18:1
TG 16:1_18:1_18:2
TG 18:0_18:0_18:1
TG 18:0_18:1_18:1
TG 18:1_18:1_18:1
TG 18:1_18:1_18:2
TG 18:1_18:2_18:2
TG 18:2_18:2_18:2
40
30
20
10
TG 14:0_16:0_18:2
TG 14:0_16:1_18:1
TG 14:0_16:1_18:2
TG 14:0_18:0_18:1
TG 14:0_18:2_18:2
TG 14:1_16:0_18:1
TG 14:1_16:1_18:0
TG 14:1_18:0_18:2
TG 14:1_18:1_18:1
TG 15:0_18:1_16:0
TG 15:0_18:1_18:1
TG 16:0_16:0_18:2
TG 16:1_16:1_16:1
TG 16:1_16:1_18:0
TG 17:0_16:0_16:1
TG 17:0_18:1_14:0
TG 17:0_18:1_16:0
TG 17:0_18:1_16:1
TG 17:0_18:1_18:1
TG 17:0_18:2_16:0
TG 18:0_18:2_18:2
TG 18:1_14:0_16:0
TG 18:1_18:1_20:4
TG 18:1_18:1_22:6
TG 18:2_18:2_20:4
2.0
1.5
1.0
0.5
D
Liver
DG 30:0 -(14:0)
DG 32:0 -(16:0)
DG 32:1 -(16:1)
DG 32:2 -(18:2)
DG 34:1 -(18:1)
DG 34:2 -(16:1)
DG 34:2 -(18:2)
DG 36:1 -(18:1)
DG 36:2 -(18:1)
DG 36:2 -(18:2)
DG 36:3 -(18:2)
DG 36:4 -(18:2)
DG 36:4 -(18:3)
DG 36:4 -(20:4)
DG 38:4 -(20:3)
DG 38:4 -(20:4)
DG 38:5 -(20:4)
DG 38:5 -(22:5)
DG 38:6 -(20:4)
DG 38:6 -(22:6)
8000
6000
4000
2000
Saturated
TG 16:0_16:0_16:0
TG 16:0_16:0_18:0
TG 17:0_16:0_18:0
TG 18:0_18:0_18:0
100
50
Unsaturated
TG 14:0_16:0_18:2
TG 14:0_16:1_18:1
TG 14:0_16:1_18:2
TG 14:0_18:0_18:1
TG 14:0_18:2_18:2
TG 14:1_16:0_18:1
TG 14:1_16:1_18:0
TG 14:1_18:0_18:2
TG 14:1_18:1_18:1
TG 15:0_18:1_16:0
TG 15:0_18:1_18:1
TG 16:0_16:0_18:1
TG 16:0_16:0_18:2
TG 16:0_16:1_18:1
TG 16:0_18:0_18:1
TG 16:0_18:1_18:1
TG 16:0_18:1_18:2
TG 16:0_18:2_18:2
TG 16:1_16:1_16:1
TG 16:1_16:1_18:0
TG 16:1_16:1_18:1
TG 16:1_18:1_18:1
TG 16:1_18:1_18:2
TG 17:0_16:0_16:1
TG 17:0_18:1_14:0
TG 17:0_18:1_16:0
TG 17:0_18:1_16:1
TG 17:0_18:1_18:1
TG 17:0_18:2_16:0
TG 18:0_18:0_18:1
TG 18:0_18:1_18:1
TG 18:0_18:2_18:2
TG 18:1_14:0_16:0
TG 18:1_18:1_18:1
TG 18:1_18:1_18:2
TG18:1_18:1_20:4
TG 18:1_18:1_22:6
TG 18:1_18:2_18:2
TG 18:2_18:2_18:2
TG 18:2_18:2_20:4
3000
2000
1000


**Figure S6**

**Supplementary Fig. 6 Lipidomic analysis of ageing and HFD challenged MLKL deficient mice.**

Wildtype (WT) and *Mlkl*$^{-/-}$ mice were fed a normal chow diet (ND) or high fat diet (HFD) for 25 weeks. Total lipid was extracted from the serum, liver and VAT and lipid species analyzed by LCMS. **(A)** Principle component analysis (PCA) plots of serum, liver and VAT of WT and *Mlkl*$^{-/-}$ mice fed a ND or HFD. (n = 8-12 biological samples from 3 independent experiments). **(B-D)** Heat maps of **(B)** ceramide abundance in the VAT grouped by carbon length, **(C)** TG in the serum and **(D)** DG and TG in the liver. Data in heat maps represent relative median abundance per group and values are normalized for median lipid content per mouse and tissue weight, as applicable. Related to **Figure 7**.

**Supplementary Table 1. qPCR primers**

| **Gene** | **Forward primer 5' – 3'** | **Reverse Primer 5' – 3'** |
|---|---|---|
| *Abcd2* | CCAACGGTTGTGGGAAAAGC | GAGACATGTATGGCCTCTGTGG |
| *Acaca* | GATGAACCATCTCCGTTGGC | GACCCAATTATGAATCGGGAGTG |
| *Acca1b* | CAGGACGTGAAGCTAAAGCCT | CTCCGAAGTTATCCCCATAGGAA |
| *Acot3* | TGCTCAGTCACCCTCAGGTAA | GCTTGGTGTTTATGCCCACG |
| *CD36* | TGTGTTTGGAGGCATTCTCA | TTTTGCACGTCAAAGATCCA |
| *Cidea* | GTGGTGGACACAGAGGAGTTC | TGGGACATACTTACTACCCGGTG |
| *Dgat1* | TGGTAGTGGGCCCAAGGTAG | TGCAGACGATGGCACCTCAG |
| *Elov6* | GAAAAGCAGTTCAACGAGAACG | AGATGCCGACCACCAAAGATA |
| *Fabp4* | AAGGTGAAGAGCATCATAACCCT | TCACGCCTTTCATAACACATTCC |
| *Fasn* | CTGACTCGGCTACTGACACG | AATGGGGTGCACAAGGAACA |
| *Il-1b* | AGTTGACGGACCCCAAAAG | AGCTGGATGCTCTCATCAGG |
| *Mlkl* | TCGATTCTCCCAACATCTTG | GGTGTAGCCTGTATAAGCCTCTG |
| *Mogat1* | TTGTCCTCGGAGGTGCAAAG | CTGGAACCAAACTGGCACCAT |
| *Nlrp3* | ATTACCCGCCCGAGAAAGG | TCGCAGCAAAGATCCACACAG |
| *Plin4* | TCAGTGGAGGAGTGTGGTCA | ACTGCCAGCTGAGCTTGTTC |
| *Ppara* | AGAGCCCCATCTGTCCTCTC | ACTGGTAGTCTGCAAAACCAAA |
| *Pparg* | GGAAGACCACTCGCATTCCTT | GTAATCAGCAACCATTGGGTCA |
| *Scd1* | CACCTGCCTCTTCGGGATTT | GGCCCATTCGTACACGTCAT |
| *Tnfa* | CCACCACGCTCTTCTGTCTA | CACTTGGTGGTTTGCTACGA |
| *Vldlr* | GAGGTCAACTGCCCTTCTCG | AGCCATCAACACAGTCTCGG |
| *18s* | GTAACCCGTTGAACCCCATT | CCATCCAATCGGTAGTAGCG |
